# Dissecting the nutritional regulations of a whole amino acid transporter family from a complex genome species: A holistic approach turning weaknesses into strengths

**DOI:** 10.1101/2024.10.11.617833

**Authors:** Soizig Le Garrec, Karine Pinel, Cécile Heraud, Akintha Ganot, Vincent Veron, Guillaume Morin, Ophélie Iorfida, Emilie Cardona, Lucie Marandel, Anne Devin, Jacques Labonne, Julien Averous, Alain Bruhat, Iban Seiliez, Florian Beaumatin

## Abstract

Amino acid transporters (AATs) are described as pivotal in maintaining circulating and cellular concentrations of AA via regulation of their expression in response to the cellular environment. Rainbow trout (RT), a complex genome species, is poorly described for AATs roles in controlling its predominant AA-based metabolism, despite representing a major challenge in the aquaculture nutrition field. Therefore, we identified the whole repertoire of AAT found in RT genome (>200), its expression in tissues and its nutritional regulations *in vitro*. Results garnered revealed the existence of different clusters of AATs, notably due to promoters bearing ATF4-related AA response elements. Moreover, the modeling of each AAT-specific cluster activities disclosed mTOR-related signaling functions of Ile and Phe, yet unknown in RT. Thus, this novel approach herein described should help to better grasp AA homeostasis in most organisms and topics such as fish nutrition and evolution.

## Introduction

In every field of biology, the concept of homeostasis is certainly one of the most important and recurrent. It could be defined as a set of vital mechanisms that enable an organism to maintain stable and optimal internal conditions for survival and normal functioning, despite recurrent changes in the external environment. Thus, the SoLute Carrier (SLC) family, which in human has over 400 members spread in more than 50 subfamilies (www.bioparadigms.org), has proved to be a key factor in maintaining homeostasis since it acts as a gatekeeper ensuring the cellular balance in ions and nutrients notably. Nonetheless, the comprehension of the role played by such family of proteins in physiological processes are still very sparse and uneven. Indeed, until recently, the SLC family was still described^1^ as one of the least studied family of genes in biology and in which 30% of the studied performed were focused only on 5% of its members, demonstrating the global lack of knowledge remaining in the field. Fairly, the reasons that explain this apparent lack of focus on this family have multiple origins, such as the fact that SLC genes encode for membrane proteins, which are difficult to express, detect and purify. In addition, each SLC subfamily is made of a great number of members, each of which being in charge of the uptake/transport of one or few substrates making even more challenging the study of individual members through functional invalidation or overexpression because of compensatory mechanisms that could be ensured by some functional SLC redundancies.

The Amino Acid Transporter (AAT) subfamily is a great example to highlight such issue. With more than 70 genes coding for AAT in human, this subfamily orchestrates the maintenance of circulating and intracellular AA pools within concentration ranges suitable for life. This was mainly exemplified by researches conducted in the medical field where some dysregulations of AAT were associated with the onset of pathologies^2^ (e.g., inherited rare diseases^3,4^, cancers^5^, type 2 diabetes^6^). Moreover, these AAT exhibit several differences notably for i) the type of AA transported (cationic - CAA, anionic - AAA and neutral - NAA), ii) the mechanism of transport (symport, uniport, antiport), iii) their ion dependencies (none or Na^+^; Na^+^/Cl^-^; H^+^; Na^+^/H^+^/Na^+^/H^+^/K^+^), iv) their cellular locations (mitochondria, lysosome, plasma membrane, golgi apparatus…) and v) their ubiquitous or tissue specific expressions^7,8^. On top of that comes the fact that AAT family is highly dynamic in response to environmental changes especially those related to nutrients^7,9–11^. Indeed, it was shown that a loss in nutritional AA availability led to the activation of the General Control Nonderepressible 2 (GCN2) kinase. Then, GCN2 phosphorylates the α subunit of eukaryotic initiation factor 2 (eIF2α) on serine 51 which represses general protein translation and upregulates the translation of the activating transcription factor 4 (ATF4). Once induced, ATF4 activates transcription of several AAT genes^8^. Thus, considering that each AAT is determined by a different combination of these specificities, trying to define the AAT-dependent rules that maintain AA homeostasis in an organism by studying each AAT independently could easily be like squaring the circle. This statement could be of great importance when knowing that it was recently suggested that SLCs operate as a coordinated network^12^ to support efficient protein translation, metabolism and physiological functions.

As previously mentioned, most of the discoveries on AATs were established in the medical field using mammalian models where most tools, protocols and methods are available. Nonetheless, we recently demonstrated that exploring AAT in diverse research areas and using unconventional models can also be highly valuable^13^. Indeed, knowledge garnered on cationic AAT (CAAT) in Rainbow Trout (RT, a carnivorous species whose metabolism is highly dependent on AA intake) has notably proven to be critical in optimizing diet formulation for fish of agronomic interest, ultimately promoting a more efficient metabolic use of the newly formulated diet and reducing its costly production, as deeply required by the Food and Agriculture Organization of the United Nations^14^. Building on this study, we now propose to extend the research to encompass all members of the AAT family in RT. To this end, the first challenge was the complexity of the RT genome, which undergone two additional whole genome duplication (WGD) events compared to mammals. As a result, we hypothesized that more than 200 different AATs may be present in the RT genome, adding to the challenge of studying the AAT family as a whole. This meant that, to overcome this challenge, it will be necessary to develop new protocols and methods that allowed the functional study of this great number of AAT in an integrated physiological cellular context and in absence of tools, especially those related to gene silencing which will be irrelevant and inefficient regarding the great redundancy of AAT in RT.

Thus, within this study, we first identified the whole AAT family present in RT genome and characterized its expression in different tissues as well as in a RT cell line. We then set-up original protocols and methods to characterize the nutrient-dependent regulations of the AAT family with a focus on AA-dependent regulations and their cellular outcomes on intracellular AA fluxes and mTOR (mechanistic Target Of Rapamycin) signaling. The overall analysis of the dataset obtained following multiple nutritional challenges to the cells enabled us to identify, for the first time in RT, the role of Ile and Phe in the activation of the mTOR pathway. Similarly, the analysis of the whole dataset proved the existence of concerted nutritional regulations of AATs expressions, in particular by those AA-dependent through the identification of ATF4-dependent regulatory sequences in their promoters, functionally validated *in vitro* and *in vivo*. Finally, the nutritional regulations of the AAT family were combined to the dataset of AA fluxes to propose an integrated model of the role played by AAT in the maintenance of cellular AA homeostasis.

## Results

### Strong conservation of sequence and expression profiles of RT AATs despite major genome duplication events in this species

Genome analysis of RT revealed 219 SLC coding genes from the AAT family (Fig. 1a). If on an average each human AAT gene (72) could be represented by 3 orthologs in RT, in fact this value greatly varied from one gene to another. Indeed, 8 human genes were not found in RT genome (EAAT4, ORNT2, GC2, PAT2, PAT3, SNAT1, SNAT5 and SFXN3) while two others (EAAT6 and 7), previously shown to have been lost in mammals but kept in ray-finned fish ^15^, were present, as 4 and 2 paralogs respectively, in RT genome. Regarding the other AAT genes, some displayed only one copy (e.g. rBAT, ATB^0,+^ or SLC38A9) or a multitude of paralogs (up to 18 when considering GAT2 paralogs). Since it was shown that part of the duplicated genes from the latest WGD are pseudogenes^16^, we analyzed their expression through RTqPCR by specifically designing primers for each AAT gene identified. From case to case, DNA sequences from paralogs were either sufficiently distinct to design paralog-specific primers or too similar to discriminate paralogs at the individual level. Therefore, the set of primers used could detect the expression of 1, 2 or 3 paralogs from the same gene as shown in Fig.1a and supplementary Table 1. Accordingly, in the pool of RT tissues analyzed (including stomach, gut, liver, muscle, ovary, spleen, kidney, brain and adipose tissue), we concluded that between 119 and 174 different AAT were expressed in adult RT organs. While looking at the theoretical protein identities of the AAT compared to human, we first noticed that AAT expressed are better conserved (around 70% identity) compared to those for which no expression could be detected (<60% identity, Fig. 1b). In the same way, we also noticed that anionic AAT (AAAT) are slightly, but significantly, more conserved compared to neutral AAT (NAAT) and CAAT (Fig. 1c). Similar conclusions have been drawn while looking at sequences of intracellular AAT compared to those located at the plasma membrane (Fig. S1a). Finally, proton-dependent AAT were those with the highest identities observed when looking at the ion dependencies (Fig. S1b) as well as were antiporters when considering transport mechanisms (Fig. S1c). Interestingly, while identities displayed by AAT were globally elevated, the subunits in charge of the trafficking of AAT in their membranes showed the lowest identity values that dropped to less than 45% on an average. Furthermore, while pursuing the comparison of the AAT family expressed in RT to its human counterpart, we observed that, despite uneven duplications events at individual levels, the proportions of the AAT were remarkably maintained whether considering the substrate transported (Fig. 1d), cellular locations (Fig. S1d), ion dependencies (Fig. S1e) and transport mechanisms (Fig. S1f). Altogether, these results highlighted a great conservation of RT AAT, when compared to human, regarding either their identities and their global expression profiles related to their specificities.

**Figure 1:**
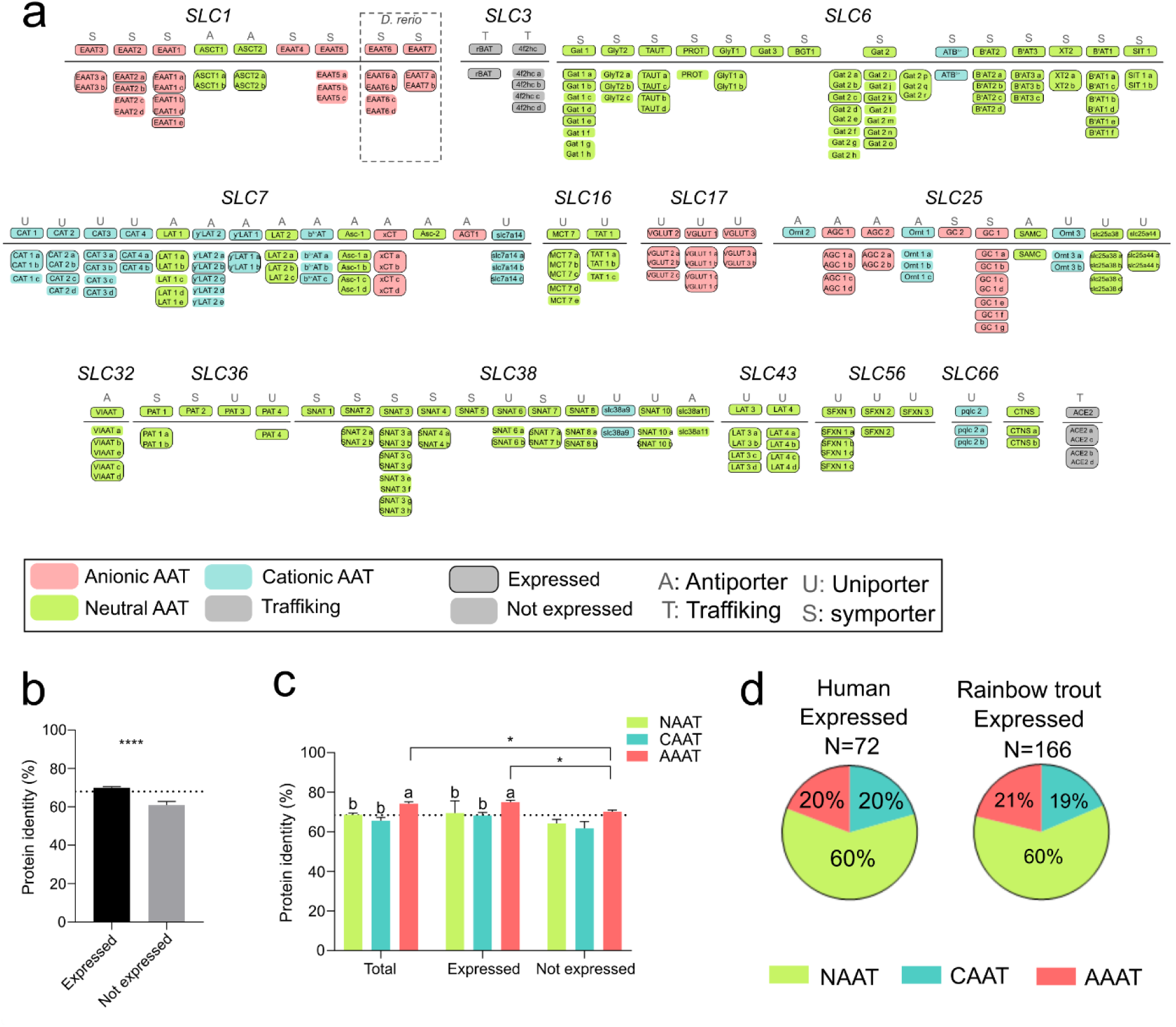
Repertoire of amino acid transporters (AAT) identified in Rainbow trout. a. AAT identified in rainbow trout genome based on known human AAT and two based on zebrafish (*D. rerio*) genome. AAT were classed according to SLC gene sub-families, categories (neutral, anionic, cationic), transport mechanism (uniporter, antiporter, symporter) and tissues expression were specified. Merged boxes indicate AAT for which the expression could not be detected individually. Gene names indicated above the grey lines are known human AAT genes while below the lines are shown their orthologs found in RT genome. b. Mean protein sequence identity, compared to human, of total AAT expressed vs. non-expressed in pool of rainbow trout tissues. c. Mean protein sequence identity by categories, compared to human, between total, expressed and non-expressed AAT. d. Proportion between each AAT category in human and rainbow trout AAT pool; Chi² proportion test between human and Rainbow trout was not significant. *: t-test statistical significance; letters: One-way Anova statistical significance. Different letters indicate a significant difference (p value <0.05).

### A global sight of the starvation-induced regulations on AAT expressions, activities and their related outcomes on mTOR signaling

We recently noticed that part of the CAAT sub-family was subjected to specific regulations orchestrated upon fluctuating nutritional conditions^13^. Nonetheless, a couple of questions remained to be elucidated. Firstly, we wondered whether such regulations are also applied to the two other sub-families (AAAT and NAAT). In other words, we sought to determine if specific cellular signatures related to AAT regulations could be defined within the whole AAT family subjected to different growing conditions. Secondly, if AAT expressions are modulated, from now on, no evidence was brought that it could be correlated to an increase in AA cellular fluxes together with their outcomes on cellular processes. Thus, taking advantage of the cell line model previously validated in our laboratory to assess the cellular aspects of nutrition in rainbow trout^13,17,18^, a set of experiments was therefore developed to address the nutritional-regulations of AAT and their outcomes on intracellular AA pools and mTOR-related signaling events known to promote anabolism through AA-dependent mechanisms. Briefly, and as shown in Fig. 2a, cells grown for 48h in control media are first subjected to different nutritional challenges for 24h (Step1). At this stage, cells are either sampled to assess AAT regulations at mRNA and protein levels or subjected to 2h starvation (Step2) to empty their AA intracellular contents. Finally, following a third step (Step3) where cells were placed back in control media, intracellular AA contents were determined through UPLC analysis as well as mTOR activation pathway *via* western blot analysis. Thus, RTH-149 cells, which shared the expression of 80% of their AAT with their original tissue (Fig. S2a), were first starved from AA and serum for 24 hours prior to assess AAT expression at transcriptional and protein levels. Therefore, we observed that 76% of the AAT expressed in RTH-149 cells are subjected to starvation-induced regulations, the majority of which represented by significant up-regulations (46%) (Fig. 2b). In the meantime, we confirmed that these regulations were specifically due to the lack of AA and serum since they were largely repealed in fed-like condition consisting of a starvation media supplemented with AA and serum (Fig. S2b). Additionally, when considering the AAT identities as well as their specificities, we noticed that only NAAT and CAAT sub-families showed significant starvation-induced up-regulation profiles (Fig. 2c and Fig. S2c). Proteomic analysis of cells subjected to similar treatments only showed a trend for NAAT up-regulations and clearly confirmed that the CAAT subfamily was highly overexpressed (Fig. 2d and Fig. S2d). Since the results gathered so far pointed to starvation-induced regulations mainly affecting CAAT and NAAT sub-families, we then investigated whether those regulations could impact the intracellular AA pools and fluxes, especially those of NAAT and CAAT since it is known that AAA pool is more controlled by metabolism than transport activities^19^. Accordingly, cells were subjected to similar nutritional regulations (Step 1) prior being shortly starved (Step 2) before to be treated back with the same complete media for 10 or 240 min (Step 3) to assess the variations of intracellular AA pools dependent on transport activities and mTOR activation levels respectively. It is important to notice that, according to previous results^18^, the step 2 starvation duration was determined to allow a significant decrease of the intracellular essential AA (EAA) pool as well as mTOR signaling but being not long enough to drive a starvation-induced regulation of AAT expression which can only be detected following a minimum of 8 hours of treatment. Consequently, we showed that NAA and CAA intracellular pools were significantly increased in an AA/Serum dependent manner (Fig. 2e and 2f) while AAA were unaffected by the nutritional past of the cells compared to those that were not exposed to a starvation (Fig. 2g). Surprisingly, we noticed that all NAA and CAA did not contribute evenly to these increases. Indeed, only 8 NAA out of 15, namely Met, Val, Ile, Phe, Ser, Gly, Cys and Gln, were responsible for the elevation of the NAA pool observed following a starvation (Fig. S2e). Furthermore, the increase in CAA pool was only supported by an increase of the intracellular Lys content while Arg and His intracellular concentrations were comparable with those of cells that did not experience a starvation (Fig S2f). When considering that the absolute intracellular concentrations of one AA greatly varied to another, the analysis of the global contribution of these fluctuations within the EAA and non-EAA (NEAA) pools revealed that cells that experienced a starvation displayed a greater ability to enrich their intracellular concentrations of EAA (Fig. 2h) while the NEAA pool was globally kept unchanged (Fig. 2i). Accordingly, this increase in intracellular EAA led to an improve activation of the mTOR pathway, exemplified by the AA/Serum dependent phosphorylation levels of the S6 protein measured (Fig. 2j). Altogether, these first results tend to demonstrate that a starvation of AA and Serum controlled the expression of a vast majority of AAT, mainly NAAT and CAAT, which in turn modulated the uptake of specific AA responsible for the activation of the mTOR pathway. Therefore, we decided to pursue our investigations on the independent roles of serum and AA in the regulation and functionalities of the AAT family.

**Figure 2:**
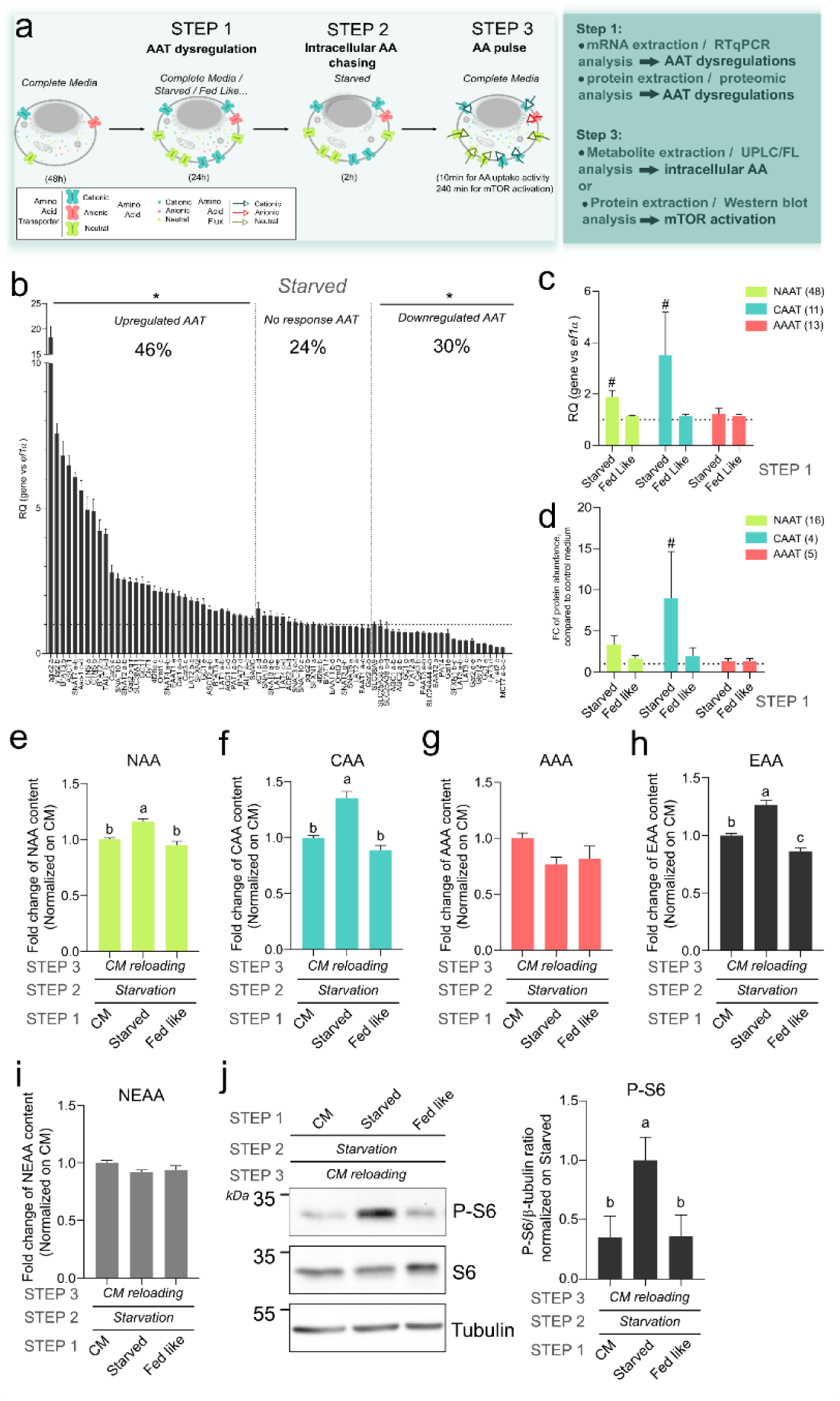
AAT dysregulation upon starvation influenced AA absorption and mTOR signalling. a. description of the experimental procedure for evaluating AAT dysregulation and effects on AA absorption and mTOR signalling. b. Relative Quantification (RQ) of AAT expressed in RTH-149 cell line in starved condition, compared to control media (CM) and normalized on EF1α expression. AAT were classed from the most upregulated to the most downregulated; percentages indicate AAT proportion in each part of the graph (N=6). c. Mean RQ for each AAT category in starved and fed-like conditions, compared to CM and normalized on EF1α expression. Numbers refer to AAT in each category (N=3). d. Mean Fold Change (FC) of protein abundance for each AAT category in starved and fed-like conditions, compared to CM. Numbers refer to AAT in each category (N=4). e.f.g.h.i FC of NAA (e), CAA (f), AAA (g), EAA (h) and NEAA (i) intracellular contents measured at the third step (N=7 for all panels except for panel g (AAA) N=3). j. Representative Western Blot and quantification of S6 phosphorylation state after CM, starved and fed-like treatments, compared to starved condition and normalized on β-tubulin quantification (N=7). # and * indicate statistical differences (T-Test, p value <0.05) compared to CM, letters: One-way Anova statistical differences. Different letters indicate a significant difference (p value <0.05).

### AA availability predominantly drive the starvation-induced AAT regulations and functionalities over serum-related responses

Following our previous observations, RTH-149 cells were starved from serum or AA to assess the regulations of AATs. We found that the response of cells starved from serum only (presence of AA in the starvation media) clearly dampened the starvation-induced regulations of AAT previously observed and shown as a red line in Fig. 3a. On the other hand, AA starvation (serum supplementation in starvation media) showed a completely different profile for which a multitude of AAT was more up-regulated when compared to the starvation condition (Fig. S3a). Considering the global regulations of each sub-family of AAT confirmed that the addition of AA in starvation media repressed the starvation-induced up-regulations of NAAT and CAAT (Fig. 3b), while serum addition had no effect on global NAAT expression but tended to reduce the starvation-induced up-regulation of CAAT sub-family (Fig. S3b). Again, AAT protein levels displayed global trends similar to those of their transcripts (Fig. 3c and S3c) where the only difference noticed was that CAAT protein levels were unaffected by serum availability (Fig. S3d-e). Moreover, intracellular AA contents measured following such nutritional challenges corroborated most of these observations since serum availability had no outcome on the increase of the starvation-induced NAA intracellular pool and a moderate negative effect on CAA’s one (Fig. S3f and S3g respectively). However, AA availability plainly abrogated the increase of CAA intracellular pool and partially reduced NAA intracellular pool (Fig. 3d-e). Finally, and consistently with the absence of global regulations of the AAAT sub-family, no significant intracellular fluctuations of the AAA pool were observed whatever the nutritional conditions considered (Fig. 3f and S3h). Interestingly, when focusing on the intracellular contents of the 9 AA previously shown for being stimulated following a starvation, we noticed that some AA levels (Fig. S3i) were controlled by both serum and AA availabilities (e.g. Lys and Met) while Val and Gly were predominantly regulated by serum and Ile, Phe, Ser, Cys and Gln by AA availability. Altogether, the regulations of the intracellular EAA pool (Fig. 3g and S3j) appeared to be controlled by AA and Serum availabilities while only serum availability led to a significant increase of the NEAA pool (Fig. 3h and S3k). Moreover, mTOR activation levels measured following those nutritional regulations partially matched with the intracellular changes in EAA observed. Indeed, phospho-S6 protein levels came across being dependent on AA availability (Fig. 3i) but not on Serum availability (Fig. S3l). Such results confirmed that the AA predominantly controlled by serum, namely Val and Gly, were not involved in the AA-dependent mTOR activation observed. Thus, all these results pointed to a predominant role of AA in the starvation-induced regulation of AAT in RT cells which globally drove the expression of NAAT and CAAT as well as the intracellular content of some specific NAA and CAA. Accordingly, we then focused our efforts in deciphering the AA-dependent regulations of the AAT family in RT.

**Figure 3:**
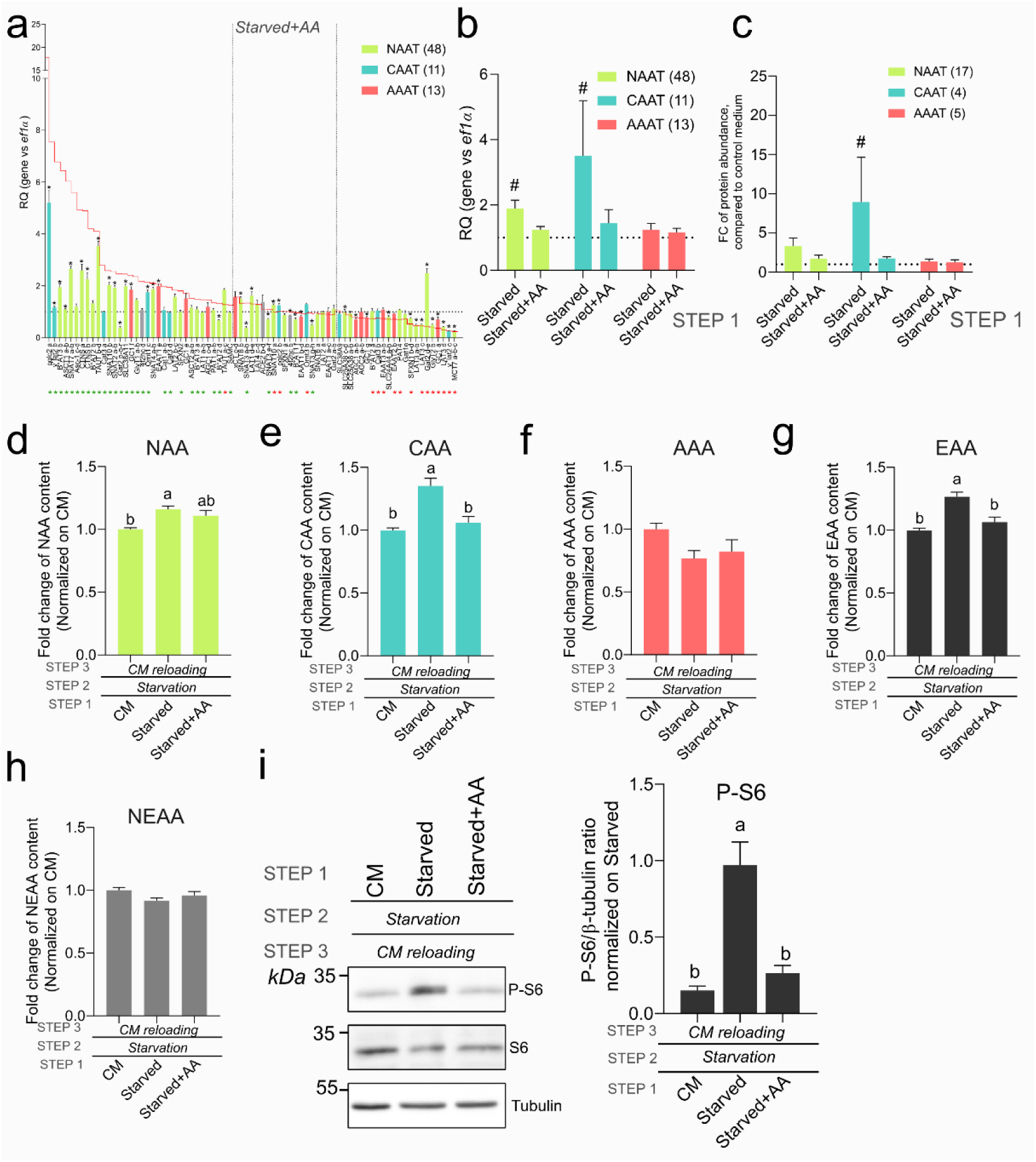
AA mainly drives AAT dysregulation influencing AA content and mTORC1 signalling. a. Relative Quantification (RQ) of AAT expressed in RTH-149 cell line in starved+AA condition, compared to CM and normalized on EF1α expression. AAT were classed from the most upregulated to the most downregulated in starved condition; red line refers to AAT profiles measured upon starved condition and colours to AAT category. Black *: t-test significant differences from CM; red and green indicated below gene names represent t-test significant differences from starved condition, depending on regulation direction (red for upregulation and green downregulation) (N=6). b.c. Mean RQ (b) and Mean Fold Change (FC) of protein abundance (c) for each AAT category in starved condition supplemented or not with AA, compared to CM and normalized on EF1α expression. #: t-test significant differences from CM. Numbers refer to AAT in each category (N=3 and N=4 respectively). d.e.f.g.h FC of NAA (d), CAA (e), AAA (f), EAA (g), NEAA (h) intracellular contents measured after step 3 and normalized to CM. Letters: One-way Anova statistical differences (N=7 for all panels except for panel f (AAA) N=3). i. Representative Western Blot and quantification of S6 phosphorylation state measured at step 3 and normalized on β-tubulin and starved condition. Letters: One-way Anova statistical differences (N=7). Different letters indicate a significant difference (p value <0.05).

### The specific contributions of EAA and NEAA in the regulations of AAT sub-families and their consequences on intracellular AA contents

Since the starvation-induced AAT regulations appeared being predominantly driven by AA availability, similar experiments were conducted while starvation media was supplemented with either EAA or NEAA (Fig. 4a and 4b). Results showed two different profiles made of AAT regulated by both EAA and NEAA (e.g. PQLC2 a) but above all some AAT specifically regulated by EAA, positively or negatively (e.g. xCT c-d or B^0^AT2 d) as well as NEAA-specific regulations (e.g. Cat1 a-b or SNAT3 a-b). Nonetheless, at global sight, the starvation-induced upregulations of NAAT and CAAT subfamilies were clearly repressed by EAA availability, while NEAA had no effect (Fig.4 c). On the other hand, at protein levels, such effects could only be confirmed for CAAT subfamily while NAAT levels were shown to be controlled by NEAA availability (Fig.4d and Fig. S4a-b). Of note, a significant increase in the global expression of the AAAT sub-family (mainly supported by xCT c-d and EAAT1 a-c upregulations) was observed at mRNA levels but such profile was not confirmed by proteomic analysis for which EAAT1 paralogs could not be detected while xCT c expression was unaffected upon each treatment. As previously, we could notice that the intracellular pools of NAA (Fig. 4e) were brought back to those of cells that did not experience a starvation, whatever the treatment considered, while CAA pool was mainly affected by EAA availability (Fig. 4f) as well as a moderate positive effect on the AAA pool (Fig. 4g). Thus, the starvation-induced increase in Met, Ile and Phe intracellular concentrations were insensitive to EAA and NEAA supplementation, while being previously described as dependent on AA (Fig. S4c), suggesting a coordinated response involving both types of AA. On the other hand, we could see that EAA availability induced a strong decrease in the intracellular content of Gly and Lys (Fig. S4c) while it increased Val intracellular concentration. Regarding NEAA availability, it appeared that the outcomes of intracellular AA fluctuations were less pronounced since it only displayed a mild repression of the starvation-induced increase in Val, Ser, Cys and Lys intracellular concentrations while strongly increasing Gln intracellular content. According to these results, the cellular pools of EAA (Fig. 4h) and NEAA (Fig. 4i) measured in these conditions showed significant differences with a considerable drop in EAA content due to EAA availability to the expense of an increase in NEAA. However, such differences, in quantity and quality, of the intracellular pools in EAA did not fit with the mTOR activation profiles in cells (Fig. 4j) since none of the phosphor-S6 protein signals measured following EAA and NEAA supplementation in the starvation media impaired the starvation-induced enhancement of mTOR activity. These results therefore suggested that the AA-sensing machinery responsible for mTOR activation in RT cells is certainly slightly different, in terms for instance of AA detected, from the one described in mammals since none of the combined profile of known activators of mTOR recapitulated the mTOR activation levels measured. On the whole, since EAA availability had more marked effects on AAT regulations and intracellular AA fluxes compared to NEAA availability, although being not neglected, we pursued our investigations on the specific role played by 4 EAA on the regulations of AAT in RTH-149 cell line.

**Figure 4:**
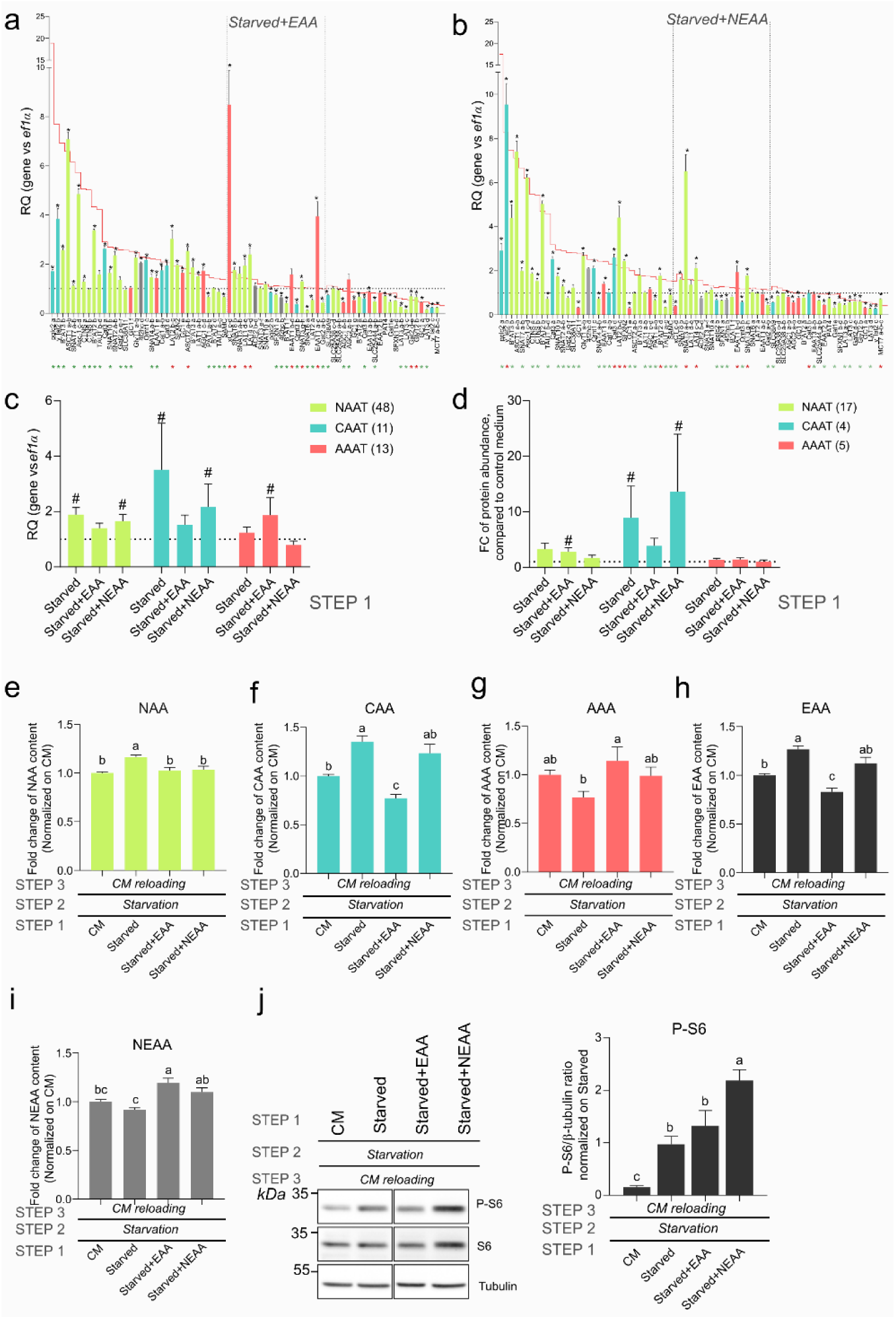
EAA mainly drove AAT dysregulation influencing AA content and mTORC1 signalling. a.b. Relative Quantification (RQ) of AAT expressed in RTH-149 cell line in starved+EAA (a) and starved+NEAA (b) conditions, normalized on EF1α expression, classed from the most upregulated to the most downregulated AAT in starved Condition. red line refers to AAT profiles measured upon starved condition and colours to AAT category. Black *: t-test significant differences from CM; red and green * indicated below gene names represent t-test significant differences from starved condition, depending on regulation direction (red for upregulation and green downregulation) (N=3). c.d. Mean RQ of gene expression (c) and Mean Fold Change (FC) of protein abundance (d) for each AAT category measured in starved condition supplemented or not with EAA or NEAA, compared to CM and normalized on EF1α expression. #: t-test significant differences from CM. Numbers refer to AAT in each category (Experiments were repeated N=3 and N=4 for panel c and d respectively). e.f.g.h.i FC of NAA (e), CAA (f), AAA (g), EAA (h), NEAA (i) intracellular contents measured at step 3 and normalized to CM. letters: One-way Anova statistical differences (N=7 for e, f, h, and i panels and N=3 for panel g=AAA). j. Representative Western Blot and quantification of S6 phosphorylation state after CM and starved with or without EAA or NEAA supplementations, compared to starved condition and normalized on β-tubulin quantification. Letters: One-way Anova statistical differences (N=7). Different letters indicate a significant difference (p value <0.05).

### Single EAA starvations differentially regulate AAT expressions, activities and signaling pathways

In our quest of dissecting the nutritional regulations of the AAT family in RT, we investigated the independent roles played by single EAA starvation by focusing on Arg, Lys, Met and Leu. Those 4 EAA were mainly picked because of their main functions in GCN2 and mTOR signaling pathways described in mammals and in fish. Moreover, Arg, Lys and Met are frequently underrepresented in plant proteins used nowadays to feed farmed fish, without really understanding the regulations induced by such restrictions. Thus, cells were incubated for 24 hours into a regular media containing all AA or deprived from one of the EAA mentioned above prior to assess the expression levels of the AAT by RTqPCR. Thus, we could observe that cells subjected to Lys or Arg starvations harbored two similar profiles for AAT expressions (Fig. 5a and 5b) where almost 50% of the AAT were up-regulated and around 30% down-regulated. Interestingly, such values were very close to the one observed upon a complete starvation (Fig. 2b) while for Met or Leu starvation (Fig. 5c and 5d respectively) very few AAT were up-regulated (20% each). On the other hand, Met starvation caused the downregulation of a very large set of AAT (48%) when Leu starvation only had a minimal effect on the down-regulation of AAT (4%). At the sight of AAT sub-families, when Arg and Lys starvations displayed a significant increase in the NAAT sub-family (Fig. 5e) and, very surprisingly, only a similar trend for CAAT sub-family, the two other starvations had absolutely no impact on the global expression of NAAT, CAAT and AAAT subfamilies. According to these observations, the AA intracellular content measured following these starvations were not, or at least very minimally, affected when considering NAA, CAA, AAA, EAA and NEAA pools (Fig. 5f to 5j respectively). It was only when considering the intracellular contents of each individual AA that slight, but significant, differences could be detected for Ile, Phe, Cys, Gln and Lys (Fig. S5a), all of which being however unaffected by Leu starvation. Altogether, despite these modest but specific changes for some intracellular AA, we could observe that only cells that undergone Arg and Lys starvation conditions could stimulate the activation of mTOR pathway (Fig. 5k) when stimulated back with a regular growing media. Consequently, we wondered whether, with the datasets of the intracellular AA contents and mTOR signaling levels gathered following all the nutritional challenges performed so far, some correlations could be established between the fluctuations of individual AA and mTOR activation in cells. Despite that Leu, Met, Arg and Lys were already shown to contribute in mTOR activation in RT cells^13,17,20,21^, only the intracellular concentrations of Ile and Phe were positively correlated with mTOR activation levels (Fig. 5l and S5b) at significant values. Therefore, we decided to evaluate whether such correlations were of biological relevance in unravelling a specific role of Ile and Phe in mTOR activation (Fig. 5m). For this purpose, cells were starved for 2h prior being treated for 4h with a starvation media supplemented or not with all EAA or the indicated AA. It turned out that indeed Ile and Phe displayed similar abilities to activate mTOR compared to its previously described AA activators. Future experiments should help to determine whether these mTOR-stimulating AA acted directly onto mTOR-dependent AA-sensing machinery or, for instance, if their uptakes involve an AAT displaying transceptor functions^22^. Overall, knowledge obtained through such single starvations indicated that i) specific cellular responses are activated depending on the identity of the missing EAA, implying that many existing AA-specific mechanisms have yet to be elucidated but also that ii) a non-negligible number of AAT regulations were commonly shared between the different starvations, suggesting the existence of a general mechanism activated following AA starvation in RT cells (Fig. S5c). Accordingly, and since this global approach revealed being powerful to identify new AA that contribute to mTOR activation in trout, we pursued the analyses of the datasets with the goal to define general rules governing the nutritional-related mechanisms of AAT expressions in RT cells.

**Figure 5:**
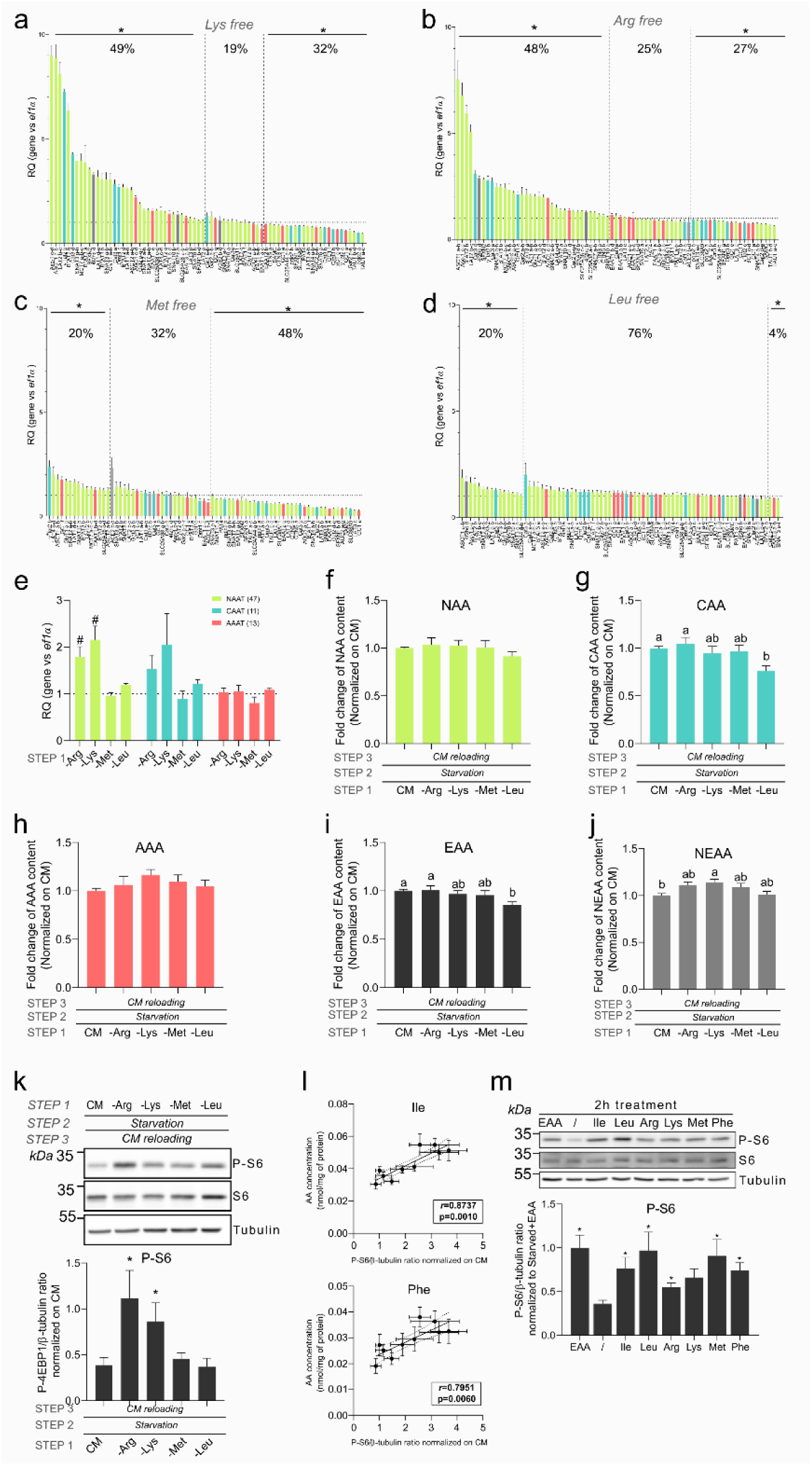
Single AA deficiencies affect AAT expression but with fewer consequences on AA absorption and mTOR signalling; while highlighting specific AA for mTOR activation. a.b.c.d. Relative Quantification (RQ) of AAT expressed in RTH-149 cell line in Lys (a), Arg (b), Met (c) and Leu (d) deficiencies, compared to CM and normalized on EF1α expression. For each treatment, AAT were classed from the most upregulated to the most downregulated AAT and colours refer to AAT category. Percentage indicate AAT proportion in each part of the graph. *: t-test significant differences from CM (N=4 for Lys and Arg, N=3 for Met and N=5 for Leu deficiencies). e. Mean RQ for each AAT category in Arg, Lys, Met and Leu deficiencies, compared to CM and normalized on EF1α expression. #: t-test significant differences from CM. Numbers refer to AAT in each category (Experiments were repeated N=4 for Lys and Arg, N=3 for Met and N=5 for Leu deficiencies). f.g.h.i.j FC of NAA (f), CAA (g), AAA (h), EAA (i), NEAA (j) intracellular contents measured at step 3 and normalized to CM. letters: One-way Anova statistical differences (N=7 for all panels except for panel h (AAA) N=3). k. Representative Western Blot and quantification of S6 phosphorylation state measured at step 3 and normalized to CM and β-tubulin. *: t-test significant differences from CM (N=7). l. Pearson correlation between isoleucine (top) and phenylalanine (bottom) intracellular contents and S6 phosphorylation state measured from the dataset of the previous 10 experimental conditions analysed. Confident intervals were shown, r=pearson coefficient; p=p-value. m. Representative Western Blot and quantification of S6 phosphorylation state after treatment with known and potential newly identify mTOR activator, compared to starved+EAA treatment and normalized on β-tubulin quantification. *: t-test significant differences from starved (/) (N=3).

### A general classification of AAT according to their nutritional-related regulations

Following the great numbers of AAT regulations observed in cells subjected to no less than 10 different nutritional challenges we first performed a hierarchical clustering of the AAT on the basis of their relative fold change expressions in each condition compared to control media (Fig. 6a). Accordingly, three different clusters, named A, B and C, could clearly be identified. The cluster A was defined as a group of AAT that were up-regulated upon starvation in an AA-dependent way while serum had a further stimulating effect on the starvation-induced up-regulations (Fig. 6b). The cluster B was mostly represented by AAT that were unaffected by nutritional conditions, while cluster C was made of AAT that were up-regulated by starvation in a serum dependent manner. Since cluster A appeared of being strongly repressed by AA availability, we assessed whether this regulation could be related to the GCN2-ATF4 pathway. First, we measured the expression levels of some specific ATF4 target genes (Fig. S6a), for which only ATF4 c-d did not display a conventional GCN2 response in most of the nutritional challenges considered. Moreover, very surprisingly but consistently with results previously obtained (Fig. 5d), Leu starvation only led to a very weak up-regulation of ASNS while other markers were left unaffected. This tended to demonstrate that RT cells do not sense Leu within the GCN2-ATF4 pathway. Overall, with the ATF4-dependent gene activation levels measured in all the nutritional conditions tested, we observed that the general expression of AAT belonging to cluster A showed significant correlations with chop, asns, ATF4 a and ATF4 b genes (Fig. 6c), suggesting a major role of the GCN2-ATF4 pathway in the regulation of this cluster. To confirm this hypothesis, cluster A was analyzed for nucleotide motif sequences commonly shared within AAT promoters from this cluster. When compared to databases, the first motif identified in cluster A almost perfectly matched with the ATF4 specific Amino Acid Response Element (AARE) motif described in mammals^23^ as an ATF4 binding site (Fig. 6d). Furthermore, such motif was found to be considerably enriched in promoters of AAT from cluster A compared to those of clusters B and C (Fig. 6e). To experimentally validated this observation, the AAT expression profiles were first assessed from cells exposed to halofuginone (Fig. S6b), a pharmacological activator of the GCN2 pathway (Fig. S6c). Again, it appeared that only the general expression of AAT from cluster A was i) specifically increased by halofuginone treatment and ii) significantly different from those of clusters B and C (Fig. 6f). To confirm the role of GCN2 in regulating AAT from cluster A, we used A92 compound, which was shown to be a a specific GCN2 inhibitor in mammals. First, we observed that the starvation-induced up-regulations of most AAT previously observed was clearly repressed upon A92 treatment (Fig. S6d) as well as those of GCN2 target genes (Fig. S6e). Then, the analysis of the AAT expressions according to the clustering clearly evidenced that A92 abrogated totally the up-regulation of cluster A while also partially repressing the expressions of cluster B and C (Fig. 6g). Altogether and since these results constituted a body of evidences demonstrating the central role of the GCN2 pathway in regulating AAT from cluster A, we finally analyzed liver samples previously collected for another study^24^. Briefly, those samples were obtained from trout fed two isoenergetic and isolipidic diets that only differed from their contents in crude protein inclusion rate (60% for the high protein diet *versus* 40% for the “low” protein diet) as well as for starch to balance the gross energy for both diets. As a result, the analysis of the AAT profiles (Fig. S6f) revealed that only AAT from cluster A were statistically more expressed in the diet providing less dietary AA compared to the high protein diet (HP) (Fig. 6h) in full accordance with the expression levels of GCN2 target genes (Fig. S6g). Beside the fact that it was the first evidence of the conservation and functionality of AARE sequences in RT, all these results confirmed the central role played by the GCN2 pathway in controlling AA homeostasis through AAT regulations in RT.

**Figure 6:**
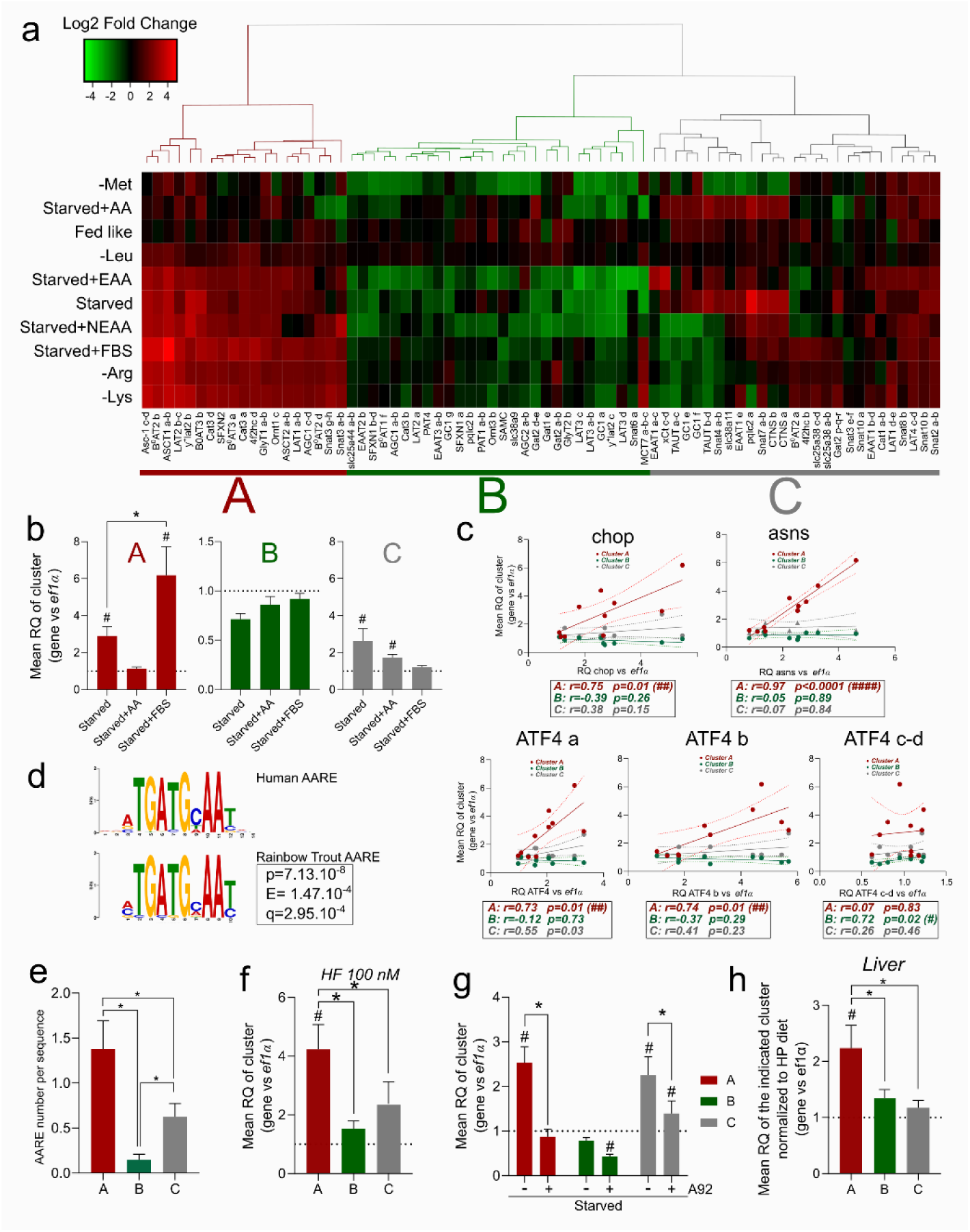
Insightful knowledge brought by clustering of AAT according to their nutritional regulations. a. Heatmap of AAT Log2 Fold change in all experimental condition tested in RTH-149 cell line. Branches indicate hierarchical clustering, allowing the identification of three clusters: A (red), B (green) and C (grey). b. Mean relative quantification (RQ) of AAT from cluster A (left), B (middle) and C (right) in the indicated condition compared to CM and normalized on EF1α expression. #: t-test significant differences from CM, *: t-test significant differences between starved conditions (N=3). c. Pearson correlation between mean RQ of each AAT cluster and mean of GCN2 target genes (chop, asns and ATF4 paralogs) in the 10 experimental conditions tested. Confident intervals were shown; pearson coefficients (r) and p-values (p) for each correlation were specified; #: indicate the level of statistical significance. d. Position Weight Matrice of discovered motif from enrichment motif analysis and comparison to Amino Acid Response Element (AARE) identified in human from JASPAR database. P = p-value of similarity; E = false positive number, q= minimum false discovery rate. e. Mean AARE number found in each AAT sequences according to cluster. *: t-test significant differences between clusters. f. Mean RQ of AAT from each cluster in CM+100nM Halofuginone condition, compared to CM and normalized on EF1α. #: t-test significant differences from CM, *: t-test significant differences between clusters (N=5). g. Mean RQ of AAT from each cluster in starved condition supplemented or not with 4µM A92, compared to CM and normalized on EF1α expression. #: t-test significant differences from CM, *: t-test significant differences between clusters (N=3). h. Mean RQ of each cluster in liver of Rainbow trout fed with High Protein (HP) or Low Protein (LP) dietary levels, compared to HP diet and normalized on EF1α expression. #: t-test significant differences from HP, *: t-test significant differences between clusters (N=9).

### Modelling the nutritional-dependent AAT activities in RT cells

Finally, while pursuing the efforts in decrypting the complex RT AAT family, we aimed to model the nutritional regulations of this family and their outcomes on AA fluxes and cellular signaling events. To do so, we first calculated the Spearman’s correlation values between each individual AA and each AAT independently, prior to proceed to a hierarchical clustering of these 1.425 values to evaluate if a specific pattern of intracellular AA fluctuations could be associated with the regulation of some specific pools of AAT. As a result, three main groups could be identified and named α, β and γ (Fig. 7a). The first thing that could be perceived was that the group α was mostly constituted of AAT from the cluster B combined to few AAT from cluster C while the group β was integrally made of AAT from cluster C and group γ was mainly represented by AAT from cluster A and few others from cluster C. In other words, while the ‘nutritional’ clustering previously carried out was based entirely on fluctuations in AAT expression observed in cells exposed to various nutrient media, it was remarkable to see that integrating the intracellular AA datasets led to virtually the same conclusions, while certainly refining them. Consequently, convinced by the robustness of this clustering, we sought to model the nutrition-dependent AA transport activities of these 3 new groups. To this end, a Bayesian approach was used to estimate the posterior values (PVs) of the effects of the three groups (α, β and γ) on intracellular variation of each AA (Fig. 7b). Thus, positive or negative PVs reflected the global import or export activities of each group and AA considered, respectively. Accordingly, this analysis tended to explain cellular AA fluxes by attributing to the group α the efflux of Gln, His, Ile, Met and Phe while it also seemed to be the major contributor for Arg uptake. Since this group is mainly constituted of AAT that do not respond to nutrients (Cluster B), it can therefore be seen as a homogenizer group in charge to equilibrate the AA intracellular pools. On the other side, the group β, only constituted of AAT repressed by serum, would be globally in charge of the efflux of Asp and Gln and the uptake of Gly, Lys and Val. Finally, the group γ, mainly constituted of AAT up-regulated by AA starvation, would be responsible for the uptake of Cys, Gln, His, Ile, Phe and Ser. For all the other AA not mentioned, since no clear differences in PVs could be estimated between groups, it was therefore suspected that these AA were shared substrates. Finally, to summarize all the findings from this study, we proposed the model of the nutritional regulations of the AAT family and their outcomes on intracellular AA contents (Fig. 7c). Thus, we demonstrated that in RTH-149 cells more than 75 AAT were expressed and could be subdivided into three clusters A, B and C according to their nutritional regulations by AA and serum. With a focus made on AA specific regulations, we could demonstrate that the cluster A was a predominant target of the GCN2 pathway notably due to an enrichment of AARE motifs in the promoters of the AAT belonging to this cluster. Moreover, the AAT have also been classified according to the modelling of their transport activities into 3 different groups (α, β and γ) each of which that could be seen as a polyvalent transporter with specificities for the uptake or efflux of key AA, among which Phe and Ile, two newly identified mTOR activators in RT.

**Figure 7:**
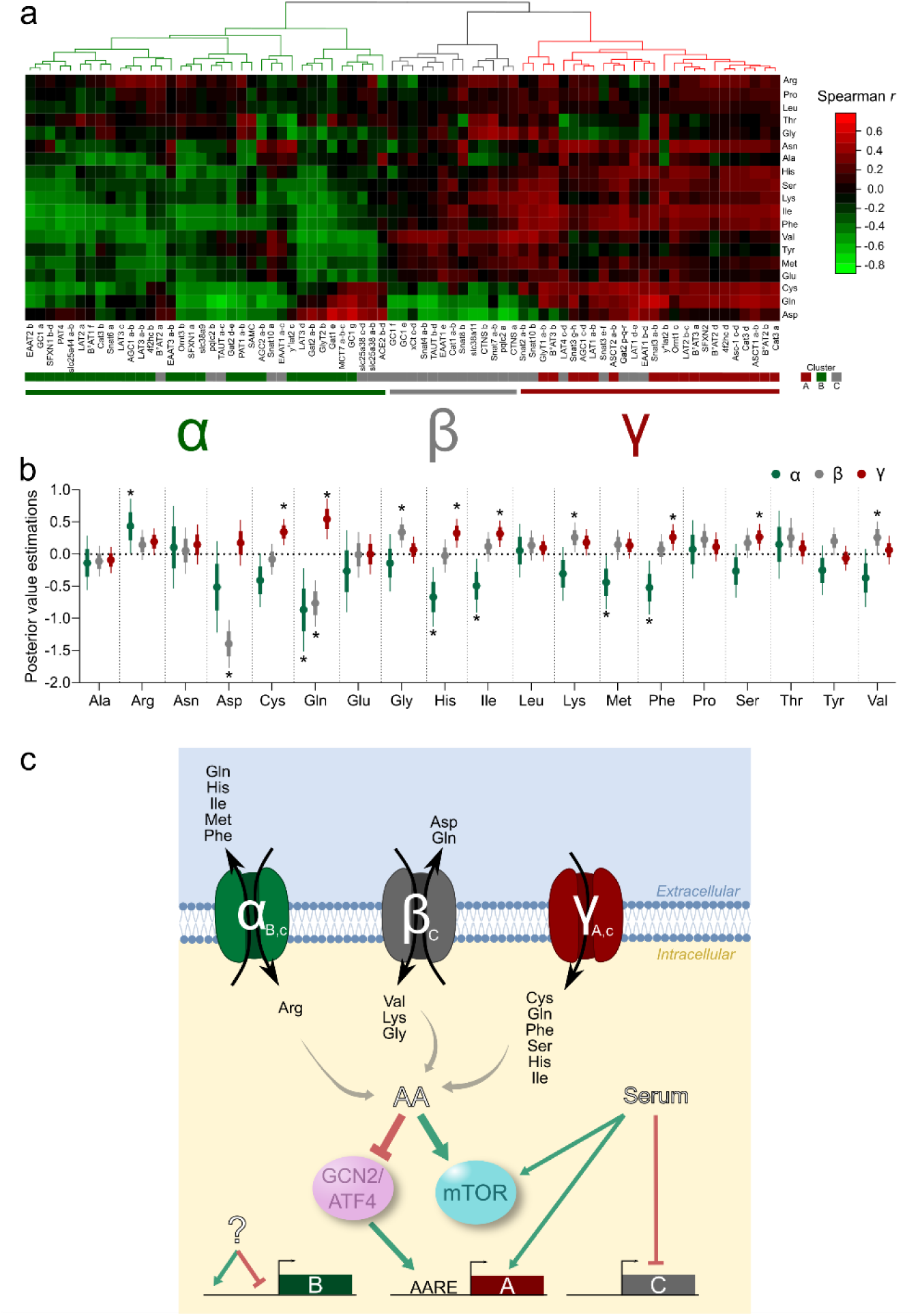
Modelling the nutritional regulations of the RT AAT family and their related transport specificities. a. Hierarchical clustering of Spearman correlation coefficient between AAT expression and AA absorption measured from the dataset of the previous 10 experimental conditions analysed. This allowed the identification of three clusters α (green), β (grey) and γ (red). Nutritional regulation of clusters identified in Fig.6 were also specified (A, B or C) b. Posterior estimation values (Pv) for each AA and each cluster activity, explaining cluster participation to AA absorption. Stars indicate that posteriors with 97.5% of distribution is different from zero. c. Proposed model of the nutritional regulations of AA fluxes for each cluster activity and the implication of GCN2/mTOR activation in these regulations. Black arrows indicate AA flux directions. Letter in each cluster referred to the specific activities estimated (α, β or γ) as well as their specific nutritional regulations outlined by hierarchical clustering (A, B or C). Green arrows: stimulating effect; crossed-out red arrows: inhibiting effect.

## Discussion

This study highlighted the great diversity and number of AAT found in RT genome as well as those expressed in its tissues and derivative cells. Results obtained demonstrated that the global profile of AAT regulations observed upon a serum and AA starvation is mainly driven by AA availability, which globally stimulated the expression of NAAT and CAAT subfamilies but not AAAT. Interestingly, we also noticed that the outcomes of these regulations on AA intracellular pools converged to impact only half of the proteinogenic AA (namely Met, Val, Ile, Phe, Ser, Gly, Cys, Gln, and Lys) where 4 are described as NEAA. This suggest specific roles played by those AA in starving cells and indicating that cells’ responses to nutrient deficiency do not spare efforts in stimulating the absorption of NEAA either, reinforcing furthermore the concept of functional AA^25^. Moreover, while looking at single AA starvation responses, our results clearly established that a combination of a general response to AA starvation and an AA-specific response are activated by cells, certainly to better cope with the starvation considered. More than that, the single starvations focusing on 4 known AA that activate mTOR showed that they are not similarly detected by the GCN2 pathway where Lys and Arg starvation induced a strong induction of GCN2 target genes while Met and Leu starvations had only a moderate, or even no impact at all, in stimulating this pathway. Such results corroborated previous observations^13^ made for which GCN2 and mTOR pathways appeared to be less interconnected in RT than in mammals which is intriguing since RT metabolism relies mainly in dietary amino acids. Moreover, the hierarchical clustering of the nutrient-dependent cellular responses of the AAT family, besides the identification of AAT regulated by the GCN2 pathway, allowed to confirmed previous statements made in mammals or in fish. First, we could see that the cluster A, mostly repressed by AA availability, is enriched in AAT localized at the plasma membrane (89%) compared to the global proportion of plasma membrane AAT expressed in RT (73%) which is consistent with an induction of the AAT from this cluster A to improve AA absorption in a context of AA restriction. Interestingly, the only two intracellular AAT found in this cluster A are mitochondrial carriers (Ornt1c and AGC1 c-d) involved in mitochondrial ornithine and aspartate/glutamate fluxes respectively, previously shown for their functions in arginine synthesis and urea cycle^26^ notably stimulated by starvation conditions. Similarly, according to WGD events that occurred in RT evolution, AAT genes were also duplicated and mainly retained in RT genome. Interestingly, for 11 (B°AT2, LAT2, y+LAT2, CAT3, 4F2hc, LAT1, AGC1, SNAT3, GC1, PQLC2 and GAT2) of the 48 human AAT orthologs expressed in RTH-149 cells (for a total of 75 AAT expressed), we could notice that subfunctionalization^27^ processes might have already occurred since paralogs from a duplicated gene were found spread into 2 different clusters (A:B, A:C or B:C). This illustrates that the AA homeostasis supported by the RT AAT family is even more complex to delineate since paralogs from a duplicated gene display very close sequence identities, so likely similar transport activities and specificities, but differences in expression levels and regulations. Therefore, in absence of extensive studies detailing all the regulations to which AAT are subjected, the functional role of some AAT will certainly be minored, if not totally ignored. Altogether, we proposed a global approach to considerably gain insight into the molecular mechanisms regulating AAT expressions and their functions in RT through the prism of nutritional regulations, exemplified and validated by the discovery that Ile and Phe are mTOR activators in RT cells. Beyond the fundamental knowledge gathered for this species of agronomic interest, this study offers to get into the complex world of AAT through an innovative and global angle to better apprehend the mechanisms involved to preserve AA homeostasis. Therefore, multiplying the use of this global approach to other stimuli, whether being related to nutrients (e.g. the responses to carbohydrate, lipids or micronutrients) or not (responses to cytokines, hormones and growth factors), will contribute to ease the global comprehension of the cellular functions supported by AAT. Moreover, since none of the protocols and methods proposed in this study required tools specifically developed for a dedicated species, this study provides a universal approach deeply required to democratize studies on the SLC family and help biology to keep moving forward in wider topics and organisms.

## Methods

### Cell culture & treatments

All in vitro experiment were conduct on RTH-149 cell line, derived from rainbow trout hepatoma (ATCC® CRL-1710, LGC standards, Molsheim, France). These cells are routinely cultured in Minimal Essential Medium (MEM, #61100-053) supplemented with 100µM Non Essential Amino Acid (NEAA, #11140-50), 1mM Sodium-pyruvate (#11360-070), 50 units/ml penicillin / 50µg/ml Streptomycin (PenStrep, #15140-122), 10% Fetal Bovine Serum (FBS, #10270-106), all provided by Gibco (Thermo Fisher scientific, Waltham, MA, USA) and 25 mM HEPES (#BP299-1, Fisher Bioreagents, Fisher scientific SAS, Illkirch Graffenstaden, France). Cells were grown at 18°C, without CO2 and at pH 7.4. Medium were change 2 to 3 times per week and cells were passaged at around 80% of confluence. Before each experiment, living cells were counted with a Cellometer K2 and AO-PI staining (#CS2-0106), both from Nexcelom Bioscience LLC (Lawrence, MA, USA). For RNA extraction, 500 000 cells were plated in 6 cm dishes, while 400 000 cells were plated in same dishes for Protein extraction. For UPLC-MS analysis, 400 000 cells were plate in 6-well plates, with 3 replicates per condition. 48h after seeding, cells were wash twice with Phosphate Buffered Saline (PBS, #2944-100, Fisher bioreagents) and treated with specific medium: Complete Medium (CM condition, constituted as mentioned above), Hank’s balanced salt solution (HBSS, #14065-056, Gibco) with 25mM HEPES was use as a starvation medium (Starved condition) and supplemented/or not with NEAA, Essential Amino Acid (EAA, #11130-036, Gibco), L-Glutamine (#25030-024, Gibco) and FBS. HBSS solution supplemented with all of these compounds were hereafter referred as Fed Like medium. For single AA starvation, a MEM medium lacking almost all AA (except Histidine, Isoleucine, Phenylalanine, Threonine, Tryptophan, Tyrosine, Valine, C4086.0500, Genaxxon Bioscience, Ulm, Germany) were used and supplemented with PenStrep, NEAA L-Glutamine, and each other AA one by one to MEM concentration, except one (minus L-Arginine (#A5006-100G), L-Lysine (#L5501-25G), L-Methionine (#M9625-25G) and L-Leucine(#L8000-50G), all provided from Sigma-Aldrich, Darmstadt, Germany). Finally, Halofuginone hydrobromide (#32481, Sigma-Aldrich), a pharmacological activator of GCN2 pathway was used at 100 nM in CM and A-92 (#Axon 2720, Axon Medchem PV, Groningen, The Netherlands), an inhibitor of GCN2 pathway was added at 4µM in Starved media. All these treatments were carried out for 24 hours before RNA extraction.

In order to assess effect of AAT regulation on intracellular AA content and mTOR activation, after these 24h treatments, cells were starved with the Starved condition during 2h, for emptying intracellular AA pool and inactivate mTOR. Cells were then reloaded with CM during 10 min before metabolites extraction and UPLC-MS analysis, or during 4h with CM lacking FBS before protein extraction for Western Blot analysis.

Finally, to assess mTOR activation level by some EAA, 48h after plating, cells were starved with HBSS and preload with NEAA for 2h, in order to inactive mTOR. Cells were then reloaded with the starvation condition containing all EAA or only one (L-isoleucine, #I5227-5G, L-phenylalanine, #P5482-25G, both provided form Sigma-Aldrich, and L-arginine, L-lysine, L-leucine, L-methionine; each at MEM concentration) or nothing during 2h, prior to protein extraction for Western Blot analysis.

### RNA extraction and quantitative RT-PCR analysis

Before RNA extraction, cells were washed with PBS solution. RNA extraction and purification were performed with RNeasy Mini Kit (Qiagen, Hilden, Germany) by using Manufacturer’s protocol. RNA concentration and integrity was assessed with a Nanodrop® ND1000 spectrophotometer.

cDNA synthesis by reverse transcriptase were carried out as previously described, as well as protocol and program for RT-qPCR analysis^13,17,18^. Briefly, cDNAs were synthetized using 1µg of RNA for each experiment, excepted for A-92 treatments where 700 ng RNA were used. All primers designed with Primer3 software and used were listed in Supplementary Table 2. Primer efficiencies were assessed using a pool of RT tissues (stomach, gut, liver, muscle, ovary, spleen, kidney, brain and adipose tissue) prior to validating amplicon size and the amplified sequence by DNA sequencing (Azenta Life Science, Leipzig, Germany). Expression of targeted genes were normalized on EF1α expression.

### Protein extraction and Western blotting

Protein extraction and Western blotting analysis were carried out as previously described^13,17,18^. Briefly, cells were lysed with RIPA buffer (#89900, Thermo Fisher scientific) supplemented with Halt protease and phosphatase inhibitor (#78442, Thermo Fisher scientific). After determination of sample protein concentration by Bicinchoninic Acid (BCA) Kit (#UP408404, Interchim, Montluçon, France) an equivalent level of protein per sample were mixed with Laemmli buffer before performed electrophoresis (12% SDS-PAGE). After protein separation and transfer, different primary antibodies were used: anti-ribosomal protein S6 (#2217L), anti-phospho-S6 (Ser235/Ser236, #4858L), and anti-β-tubulin (#2146S), all provided from Cell Signaling Technologies (Danvers, MA, USA). Chemiluminescent revelations were performed with iBright 1500 imager (Thermo Fisher Scientific) and quantification were assessed thanks to ImageJ Software (v1.53k, Java 1.8.0_172) and by using β-tubulin for normalization.

### Proteomic analysis

For proteomic analysis, proteins were extracted exactly as described above. Proteomic analysis were carried out by the ProteoToul Services – Proteomics Facility of Toulouse, France, as described previously^28^. Briefly, after preparation and digestion, samples were analyzed thanks to nano-liquid chromatography (LC) associated to tandem mass spectrometry: an UltiMate 3000 system (NCS-3500RS Nano/Cap System; ThermoFisher Scientific) and an Orbitrap Exploris 480 mass spectrometer supplied with a FAIMS Pro device (ThermoFisher Scientific). Peptide sequences were assigned against rainbow trout records in the Uniprot protein database and protein abundance of peptide identified as AAT were extracted and analyzed.

### Amino acid extraction and UPLC analysis

For analyzing AAT regulation effects on AA absorption, intracellular AA content were assessed by UPLC analysis, exactly as previously described^17^. Briefly, after 3 washes with ice-cold PBS, polar metabolites were extracted with an MeOH/H2O mix (80%/20%). Samples derivatization were performed with an AccQTag kit (#186003836, Waters) by following manufacturer’s instruction. AA content analysis were done thanks to an Acquity H-Class PLUS (Waters, Saint-Quentin-en-Yvelines, France) Alliance System (2695 separation module) associated to a Acquity UPLC Fluo Detector (Waters). All proteinogenic AA content (except Tryphophan) were identified thanks retention time compared to standards. AA content was then normalized on total protein amount, assessed with BCA Kit.

### Liver samples

For assessing AAT expression profile *in vivo*, RNA extracted from liver samples were taken from a previous study^24^. Briefly, after 4 days of total starvation, juvenile rainbow trout were fed during 4 days with two different experimental diet characterized by different ratios between carbohydrates and proteins. The first diet contained a high protein level (∼60% of crude protein, and low carbohydrate level, called “High protein” diet, HP) while the second diet contained a lower protein level (∼40% of crude protein, higher carbohydrate level allowed by inclusion of gelatinized starch, called “Low protein” diet, LP). Livers were sampled 6h after the last meal.

### Bioinformatics analysis

#### AAT identification

AAT were identified in rainbow trout genome thanks to genome annotation on NCBI database (OmykA_1.1 genome, https://www.ncbi.nlm.nih.gov/gdv) and by blasting sequences against human AAT using NCBI BLAST (Basic Local Alignment Search Tool, https://blast.ncbi.nlm.nih.gov/Blast.cgi). This was also complemented by blasting human AAT sequence against rainbow trout genome to ensure that unannotated genes were not overlooked.

#### AAT Hierarchical clustering

Clustering of AAT according to their nutritional regulation were performed using R Studio software (v4.3.0) by clustering Log2 Fold Change (FC) mRNA expression in each experimental condition test (Starved +/− nutrients and single AA deficiencies) with the Ward method.

#### Motif enrichment analysis

Based on AAT belonging to cluster A, a motif enrichment analysis was conducted with MEME Suite^29,30^ (Multiple Em for Motif Elicitation, v5.5.5). DNA sequences of more or less 1000bp from the transcription-starting site of Cluster A AAT were loaded and Classic Mode were used for motif discovery. Sequences sites distribution were assessed with One Occurrence Per Sequence (oops) setting, for identifying a maximum of 10 enriched motifs from 6 to 15 bp. To assign an enriched motif to a transcription factor binding site, the TOMTOM tool (Motif Comparison Tool^31^) were used, comparing the newly discovered motif with databases of known motif (e.g Jaspar, Uniprobe, jolma…). To identify all matching sites of the newly discovered motif, a Simple Enrichment Analysis (SEA^32^) were conducted for each cluster by taking DNA sequences of more or less 1000bp from the transcription-starting site.

#### Correlations

Correlations between AAT and AA content in all experimental conditions were conducted with R Studio (v4.3.0). Non-parametric correlation coefficients (Spearman correlation) were then clustered with the method previously described.

#### Description of the statistical model

In order to understand the dynamics of overall AA uptake by AAT, we built a model to infer the effects of AAT expression on intracellular AA content. AAT expression and AA content were Log2 fold change (FC) of mRNA expression of each AAT and log2 FC of AA content, both normalized on control condition (CM). To simplify our model and allow better parameters estimation, AAT were grouped according to groups previously determined, α, β and γ. We thus calculated the mean expression of AAT per treatment and per cluster (T_j,k_) to create k = 3 explicative factors for each of the j experimental conditions. For this model, we assumed that intracellular content of each of the i AA follow a Gaussian distribution with a mean µ_i,j_ and variance 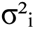 as follows:

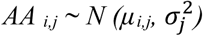

With

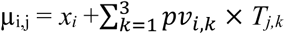

Where:

x_i_: the base value of AA uptake when all AAT expression did not vary
Pv_i,k_: the parameter effect of AAT expression on the i^th^ AA and k^th^ group.
T_j,k_: the explicative factor value for the j^th^ experimental condition and the k^th^ group.

This model assumes that 1/ AA content variance depends on experimental conditions, as demonstrated in experiments described above, 2/ variance in uptake differs between each AA and 3/ AA content might not be equal to zero even without AAT expression variation, assuming a basal AA flux.

The model was created with R Studio software (v4.3.0) using rjags and MCMCvis packages. Parameter estimation was performed using Bayesian inference framework. Relatively non-informative priors were chosen for μ_i,j_, 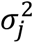, x_i_ and Pv parameters, in order to ensure rapid convergence (See https://doi.org/10.57745/WZL39N for model code, priors and data). The Pv, i.e effect of AAT groups on uptake of a given AA, x_i_ and 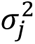 were estimated with 3 independent MCMC (Markov chain Monte Carlo) samplings with 10000 iterations each.

Model fit was assessed by calculating the percentage of explained variation in AA concentration. Posteriors parameters, named Pv, were analyzed for each AA and AAT groups, with Pv = 0 meaning that AAT group don’t have any effect on AA absorption, Pv > 0 that AAT groups have a positive effect and Pv < 0 a negative one. In practice, we considered a parameter to be statistically different from zero when 97.5% of its density probability was above or below zero.

### Statistical analysis

All data were assessed for distribution normality by Shapiro-Wilk test, and mean comparisons were assessed by two-tailed parametric T test or non-parametric U test for two-by-two comparisons, or by two-tailed parametric one-way Anova with Tuckey’s post-hoc test or non-parametric Kruskal-Wallis test and Dunn comparison test for multiple comparisons. All error bars correspond to Standard Error of the Mean (sem). Proportion analysis were assessed with Chi2 test, while correlations between AA and mTOR activation and between GCN2 target genes and global cluster mRNA expression were assessed with Pearson correlations. The significance threshold was always positioned at p<0.05.

## Data and code availability

The raw data and the code developed for this study are available on the french “*Recherche Data Gouv* multidisciplinary repository » following this link: https://doi.org/10.57745/WZL39N

## Acknowledgements

We deeply acknowledge the financial supports received from ANR JCJC (grant number ANR19-CE20-0003-01), “Université de Pau et des Pays de l’Adour” (UPPA-2018-01), the “Département INRAE de Physiologie Animale et Système d’élevage” (PhD cofounding) as well as the Aquaexcel3.0 project from Horizon Europe (871108).

## Author contributions

Conceptualization: F.B. and I.S.; methodology: S.L.G., K.P, C.H., A.D., J.A. A.B., J.L and F.B.; software: R Studio software (v4.3.0), ImageJ Software (v1.53k, Java 1.8.0_172); formal analysis: S.L.G., K.P, C.H., A.G., E.C., L.M., J.L, O.I., V.V. and G.M.; writing: S.L-G. and F.B.; editing: all authors; supervision: F.B. and I.S.; project administration: F.B. and I.S.; and funding acquisition: F.B.

## Competing interests

The authors declare no conflict of interest.

## supplementary figures and legends

**Figure S1:**
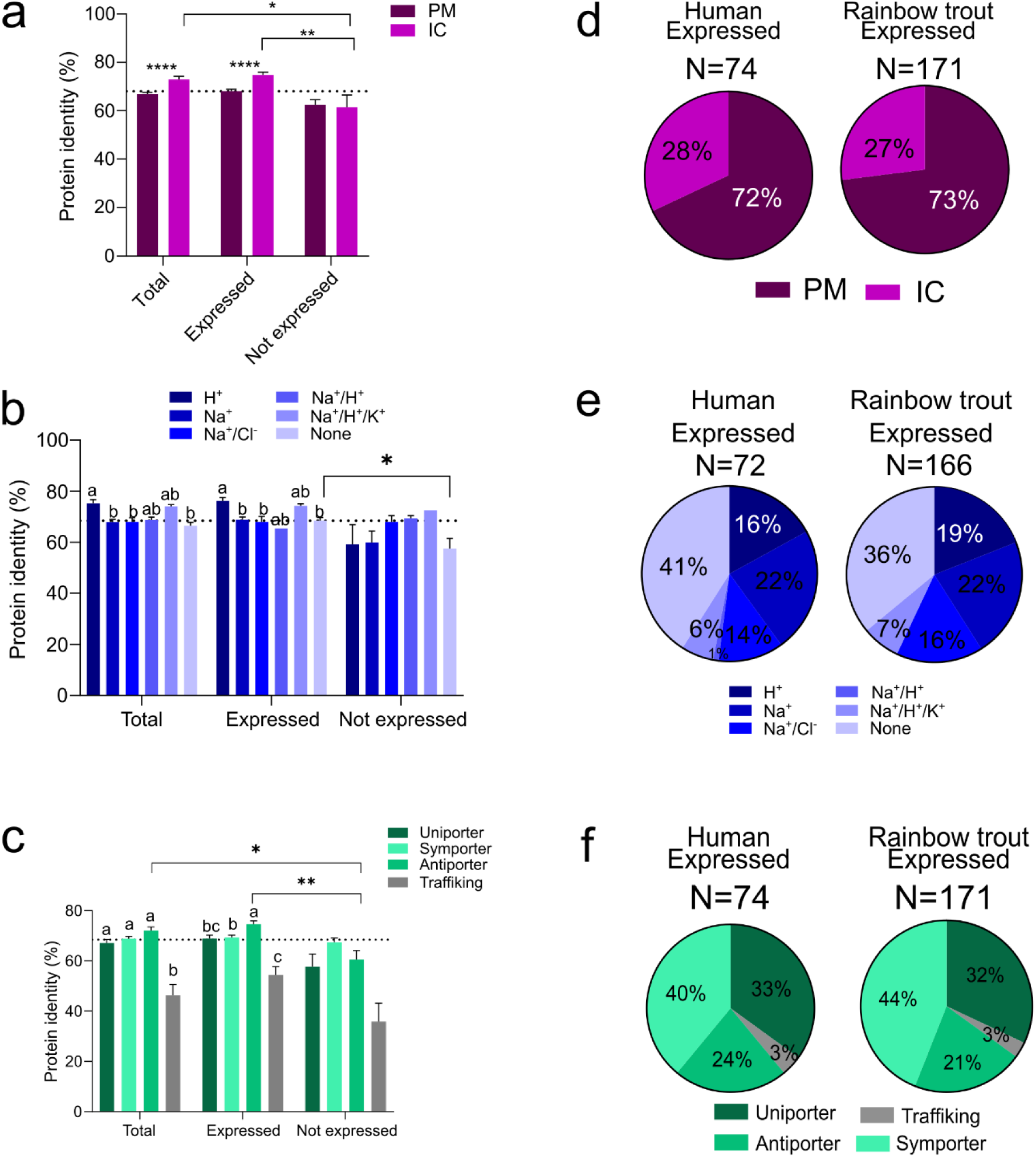
Repertoire of amino acid transporters (AAT) identified in Rainbow trout. a.b.c Mean protein sequence identity, compared to human, by (a) cellular location (PM: plasma membrane, IC: intracellular), (b) ion dependency and (c) transport mechanism between total, expressed and non-expressed AAT. d.e.f Proportion between AAT according to (d) cellular location, (e) ion dependency and (e) transport mechanism in human and rainbow trout AAT pool; Chi² proportion test between human and Rainbow trout was not significant. *: t-test statistical significance; letters: One-way Anova statistical differences. Different letters indicate a significant difference (p value <0.05).

**Figure S2:**
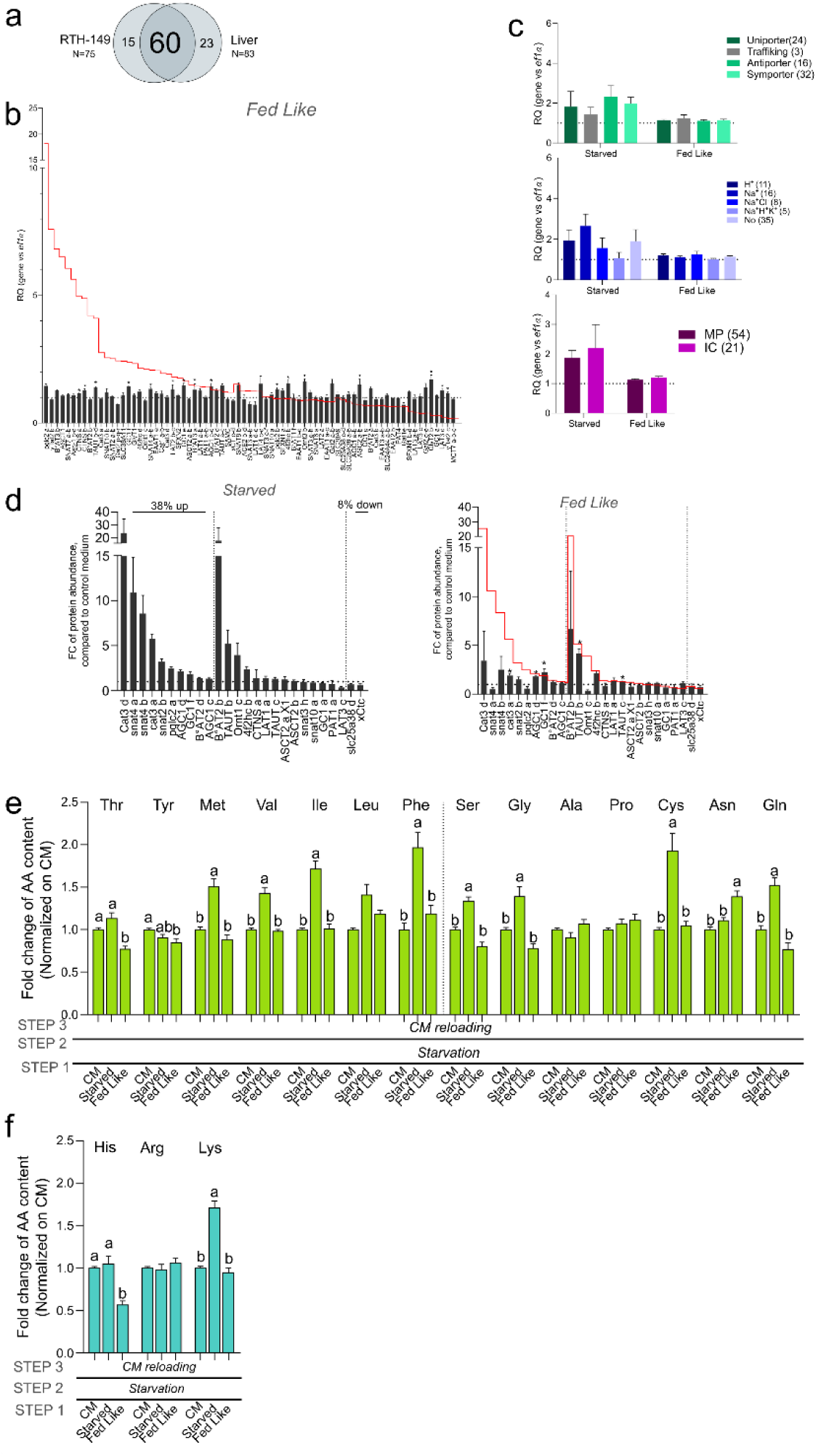
AAT dysregulation in starvation influence AA absorption and mTOR signalling. a. Venn diagram of AAT expressed in liver and in RTH-149 cell line. b. Relative quantification (RQ) of AAT in fed-like condition, compared to CM and normalized on EF1α expression, classed from the most upregulated to the most downregulated AAT in starved condition. Red line refers to AAT profiles measured upon starved conditions (N=3). C. Mean RQ for AAT groups according to transport mechanism, ion dependency and cellular location in starved and fed-like conditions, compared to CM and normalized on EF1α expression. Numbers refer to AAT in each category (N=3). d. FC of protein abundance in starved (left panel) and fed-like (right panel) conditions, compared to CM, of AAT identified in proteomic analysis, classed from the most upregulated to the most downregulated in starved conditions. Red line refers to starved level (N=4). e.f. FC of individual NAA (e) and CAA (f) intracellular contents measured after step 3 and normalized to CM. Letters: One-way Anova statistical differences (N=7). Different letters indicate a significant difference (p value <0.05).

**Figure S3:**
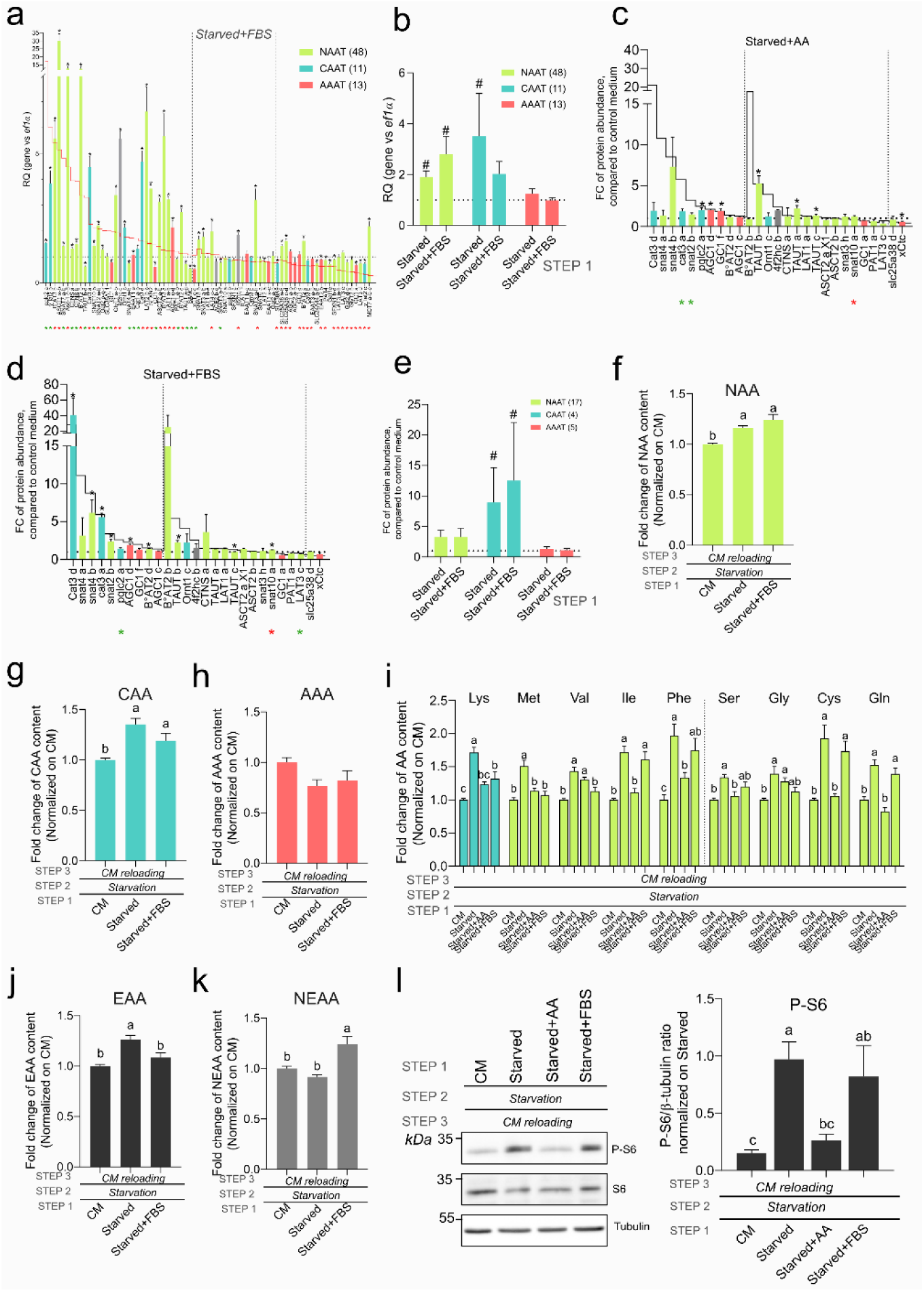
AA mainly drove AAT dysregulation influencing AA content and mTORC1 signalling. a. Relative Quantification (RQ) of AAT expressed in RTH-149 cell line in starved+FBS condition, compared to CM and normalized on EF1α expression. AAT were classed from the most upregulated to the most downregulated in starved condition; red line refers to AAT profiles measured upon starved condition and colours to AAT category. Black *: t-test significant differences from CM; red and green indicated below gene names represent t-test significant differences from starved condition, depending on regulation direction (red for upregulation and green downregulation) (N=3). b. Mean RQ for each AAT category in starved condition supplemented or not with FBS, compared to CM and normalized on EF1α expression. #: t-test significant differences from CM. Numbers refer to AAT in each category (N=3). c.d. FC of protein abundance in starved+AA (c) and starved+FBS (d) conditions, compared to CM, of AAT identified in proteomic analysis, classed from the most upregulated to the most downregulated AAT in starved condition. Black line refers to starved level, colours to AAT category. *: t-test significant differences from CM (N=4). red and green * indicated below gene names represent t-test significant differences from starved condition, depending on regulation direction (red for upregulation and green downregulation). e. Mean Fold Change of protein abundance for each AAT category in starved condition supplemented or not with FBS, compared to CM and normalized on EF1α expression. #: t-test significant differences from CM. Numbers refer to AAT in each category (N=4). f.g.h. FC of NAA (f), CAA (g), AAA (h), intracellular contents measured after step 3 and normalized on CM. Letters: One-way Anova statistical differences (N=7 for all panels except for panel h (AAA) N=3). i. FC of the 9 AA identified as being subjected to intracellular enrichment following a starvation. Intracellular contents of the indicated AA were measured at step 3 and normalized on CM. Letters: One-way Anova statistical differences (N=7). j.k. FC of EAA (j), NEAA (k) intracellular contents measured after step 3 and normalized on CM. Letters: One-way Anova statistical differences (N=7). l. Representative Western Blot and quantification of S6 phosphorylation state measured at step 3 and normalized on β-tubulin and starved condition. Letters: One-way Anova statistical differences (N=7). Different letters indicate a significant difference (p value <0.05).

**Figure S4:**
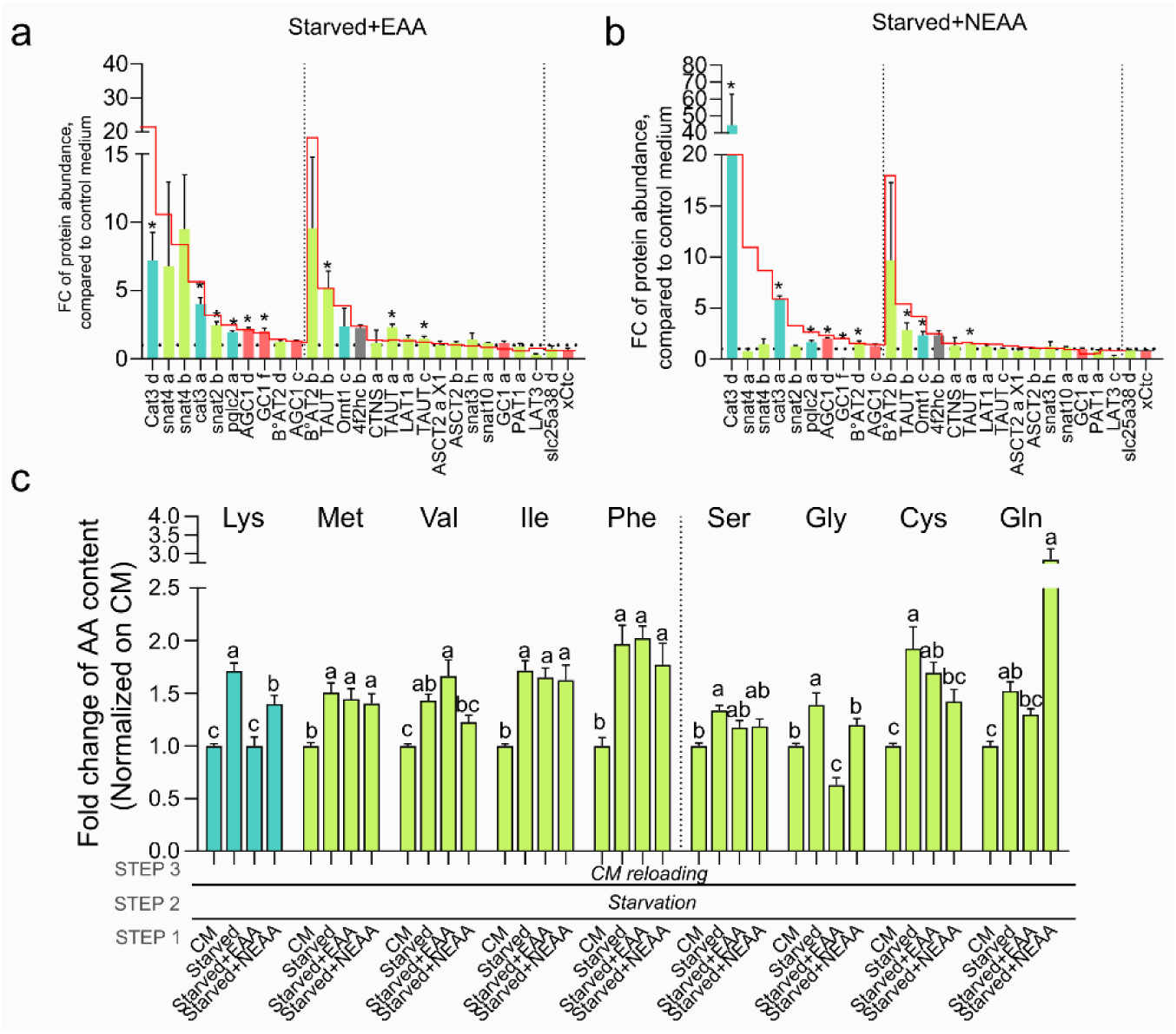
EAA mainly drove AAT dysregulation influencing AA content and mTORC1 signalling. a.b. FC of protein abundance in starved+EAA (b) and starved+NEAA (d) conditions, compared to CM, of AAT identified in proteomic analysis, classed from the most upregulated to the most downregulated AAT in starved condition. Red line refers to starved level, colours to AAT category. *: t-test significant differences from CM (N=4). c. c. FC of the 9 AA identified as being subjected to intracellular enrichment following a starvation. Intracellular contents of the indicated AA were measured at step 3 and normalized to CM. Letters: One-way Anova statistical differences (N=7). Different letters indicate a significant difference (p value <0.05).

**Figure S5:**
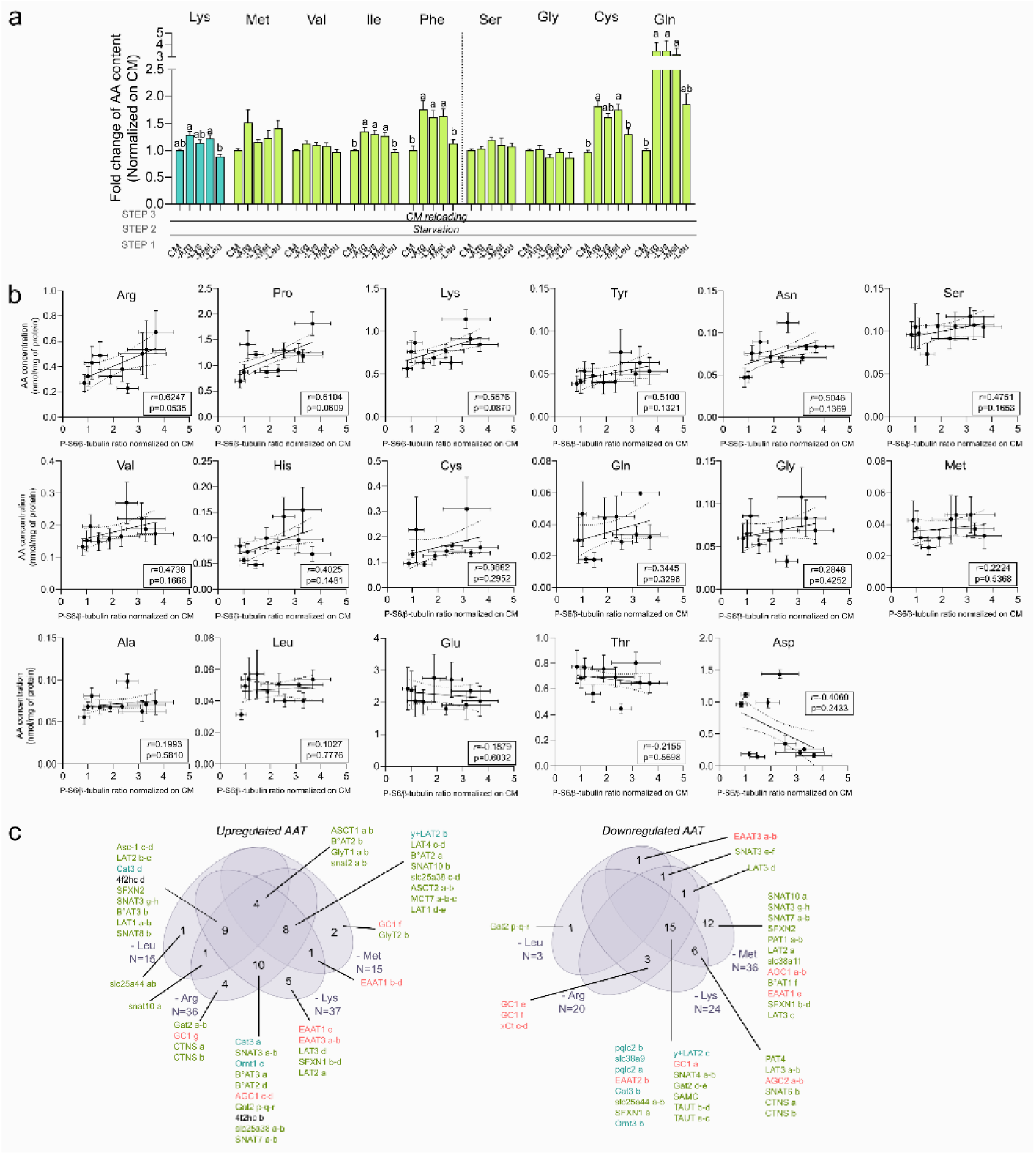
Single AA deficiencies affect AAT expression but with fewer consequences on AA absorption and mTOR signalling; while highlighting specific AA for mTOR activation. a. FC of the 9 AA identified as being subjected to intracellular enrichment following a starvation. Intracellular contents of the indicated AA were measured at step 3 and normalized to CM. letters: One-way Anova statistical differences (N=7). b. Pearson correlation between the indicated AA intracellular contents and S6 phosphorylation state measured from the dataset of the previous 10 experimental conditions analysed. Confident intervals were shown, r=pearson coefficient; p=p-value. c. Venn diagram of upregulated (left) and downregulated (right) AAT in each single deficiencies, colours refer to AAT category.

**Figure S6:**
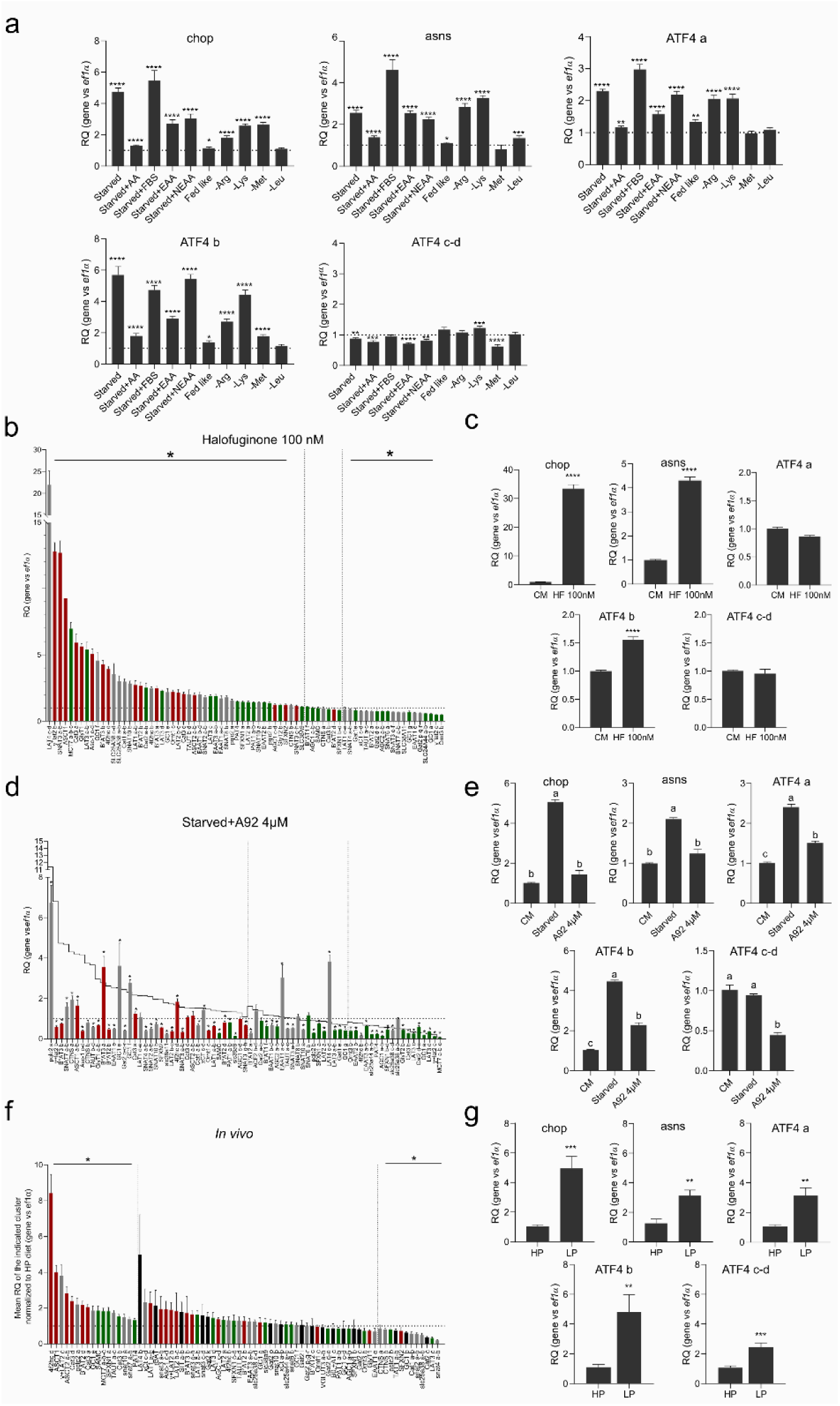
Insightful knowledge brought by clustering of AAT according to their nutritional regulations. a. Relative Quantification (RQ) of GCN2 target genes (chop, asns and ATF4 paralogs) in RTH-149 cell line in the 10 experimental conditions tested, compared to CM and normalized on EF1α expression. *: t-test significant differences level compared to CM. b. RQ of AAT in CM+100nM Halofuginone condition, compared to CM and normalized on EF1α expression, classed from the most upregulated to the most downregulated AAT. Colours refer to cluster. *: t-test statistical differences from CM (N=5). c. RQ of GCN2 target genes in CM+100nM Halofuginone, compared to CM and normalized on EF1α expression. *: t-test significant differences level compared to CM (N=5). d. RQ of AAT in starved+A92 conditions, compared to CM and normalized on EF1α expression, classed from the most upregulated to the most downregulated AAT in starved condition. Grey line refers to AAT profiles measured upon starved condition and colours to clusters. *: t-test statistical differences from CM (N=3). e. RQ of GCN2 target genes in starved condition supplemented or not with 4µM A92, compared to CM and normalized on EF1α expression. Letters: One-way Anova statistical differences. (N=3). f. RQ of AAT *in vivo* in liver of rainbow trout fed with Low Protein (LP) diet, compared to High Protein diet (HP) and normalized on EF1α expression, classed form the most upregulated to the most downregulated. Colours refer to clusters; black bars correspond to AAT only express *in vivo,* so not found in RTH-149 cells. *: t-test statistical differences from CM (N=9). g. RQ of GCN2 target genes *in vivo* in liver of rainbow trout fed with LP and HP diets, compared to HP and normalized on EF1α expression. *: t-test significant differences level compared to HP (N=9).

## Notes

https://doi.org/10.57745/WZL39N

### Competing Interest Statement

The authors have declared no competing interest.

