## Supplementary Table 1 for "Dissecting the nutritional regulations of a whole amino acid transporter family from a complex genome species: A holistic approach turning weaknesses into strengths"

**Supplementary Table 1: Rainbow trout AAT identification and expression profile.**

*In silico* identification of rainbow trout AAT, compared to human, with paralog numbers, gene ID and attributed names. Blue italic AAT refer to genes identified compared to *D. rerio* AAT. Expression profiles were assessed in a pool of RT tissues (stomach, gut, liver, muscle, ovary, spleen, kidney, brain and adipose tissue) or specifically in liver or in RTH-149 cells as indicated. Y (Yes) and N (No) means that an expression for the indicated paralog was detected or not, respectively. Substrate categories A, N, C or T : Anionic, Neutral, Cationic or Trafficking sub-unit repsectively

| Human transporter | Human gene | Paralog Numbers | Substrate Category | Chromosom | geneID | Attributed name | *In vivo* expression | Liver expression | RTH-149 expression |
|---|---|---|---|---|---|---|---|---|---|
| EAAT3 | slc1a1 | 2 | A | 9 | 110531767 | *EAAT3 a* | Y | Y | Y |
| | | | | 20 | 110500880 | *EAAT3 b* | Y | Y | Y |
| EEAT2 | slc1a2 | 4 | A | 1 | 110522323 | *EAAT2 a* | Y | N | N |
| | | | | 2 | 110497158 | *EAAT2 b* | Y | N | Y |
| | | | | 26 | 118944446 | *EAAT2 d* | N | N | N |
| | | | | 6 | 110527005 | *EAAT2 c* | N | N | N |
| EEAT1 | slc1a3 | 5 | A | 1 | 110502566 | *EAAT1 a* | Y | N | Y |
| | | | | 5,2 | 110524216 | *EAAT1 c* | Y | N | Y |
| | | | | 5,1 | 110523776 | *EAAT1 b* | Y | Y | Y |
| | | | | 12 | 110537147 | *EAAT1 d* | Y | Y | Y |
| | | | | 6 | 110525225 | *EAAT1 e* | Y | Y | Y |
| ASCT1 | slc1a4 | 2 | N | 1 | 110531632 | *ASCT1 a* | Y | Y | Y |
| | | | | 23 | 110531639 | *ASCT1 b* | Y | Y | Y |
| ASCT2 | slc1a5 | 2 | N | 24 | 110503495 | *ASCT2 a* | Y | Y | Y |
| | | | | 27 | 110507436 | *ASCT2 b* | Y | Y | Y |
| EAAT4 | slc1a6 | 0 | A | Not Found | | | | | |
| EAAT5 | slc1a7 | 3 | A | 5 | 110524687 | *EAAT5 a* | N | N | N |
| | | | | 8 | 110531019 | *EAAT5 b* | N | N | N |
| | | | | 28 | 110509189 | *EAAT5 c* | N | N | N |
| *EAAT6* | *slc1a8* | *4* | *A* | *5* | *110524108* | *EAAT6 b* | *Y* | *N* | *N* |
| | | | | *1* | *110489874* | *EAAT6 a* | *Y* | *N* | *N* |
| | | | | *18* | *110497072* | *EAAT6 d* | *N* | *N* | *N* |
| | | | | *7* | *110497076* | *EAAT6 c* | *N* | *N* | *N* |
| *EAAT7* | *slc1a9* | *2* | *A* | *13* | *110486646* | *EAAT2 a* | *Y* | *Y* | *N* |

| | | | | | | | | | |
|---|---|---|---|---|---|---|---|---|---|
| [illegible] | [illegible] | [illegible] | [illegible] | *12* | *110538991* | *EAAT2 b* | *Y* | *Y* | *N* |
| rBAT | slc3a1 | 1 | T | 20 | 110499302 | *rBat* | Y | Y | N |
| 4f2hc | slc3a2 | 4 | T | 10 | 110533969 | *4f2hc a* | N | N | N |
| | | | | 21 | 110500797 | *4f2hc b* | Y | Y | Y |
| | | | | 27 | 110507427 | *4f2hc c* | N | N | N |
| | | | | Y | 110509955 | *4f2hc d* | Y | Y | Y |
| GAT1 | slc6a2 | 8 | N | 7 | 110528658 | *Gat1 b* | Y | Y | N |
| | | | | 17 | 110504503 | *Gat1 d* | Y | N | N |
| | | | | 9 | 110531998 | *Gat1 a* | N | N | N |
| | | | | 16 | 110492407 | *Gat1 c* | Y | N | N |
| | | | | 15 | 110499848 | *Gat1 f* | N | N | N |
| | | | | 18 | 110496877 | *Gat1 e* | Y | Y | Y |
| | | | | 15 | 110499842 | *Gat1 g* | N | N | N |
| | | | | 21 | 110514342 | *Gat1 h* | N | N | N |
| GlyT2 | slc6a5 | 3 | N | 6 | 110526924 | *GlyT2 b* | Y | N | Y |
| | | | | 2 | 110499833 | *GlyT2 a* | N | N | N |
| | | | | 26 | 110499833 | *GlyT2 d* | N | N | N |
| TAUT | slc6a6 | 4 | N | 7 | 110528591 | *TAUT a* | Y | Y | Y |
| | | | | 17 | 110493294 | *TAUT c* | Y | Y | Y |
| | | | | 9 | 110531953 | *TAUT b* | Y | Y | Y |
| | | | | 16 | 110492485 | *TAUT d* | Y | Y | Y |
| PROT | slc6a7 | 1 | N | 10 | 110533742 | *PROT* | N | N | N |
| GlyT1 | slc6a9 | 2 | N | 8 | 110530321 | *GlyT1 a* | Y | Y | Y |
| | | | | 28 | 110508726 | *GlyT1 b* | Y | Y | Y |
| GAT3 | slc6a11 | 0 | N | Not Found | | | | | |
| BGT1 | slc6a12 | 0 | N | Not Found | | | | | |
| | | | | 2 | 110493383 | *Gat2 a* | Y | N | Y |
| | | | | 30 | 110521389 | *Gat2 b* | Y | N | Y |
| | | | | 15 | 110490150 | *Gat2 c* | Y | Y | N |
| | | | | 15 | 110499854 | *Gat2 d* | Y | Y | Y |
| | | | | 21 | 110499843 | *Gat2 e* | Y | Y | Y |
| | | | | 27 | 110507993 | *Gat2 f* | N | N | N |
| | | | | 7 | 110528327 | *Gat2 g* | N | N | N |

| | | | | | | | | | |
|---|---|---|---|---|---|---|---|---|---|
| GAT2 | slc6a13 | 18 | N | 7 | 110528327 | *Gat2 h* | N | N | N |
| | | | | 7 | 110527587 | *Gat2 i* | Y | Y | N |
| | | | | 7 | 110528308 | *Gat2 j* | N | N | N |
| | | | | 7 | 110528307 | *Gat2 k* | Brain | Y | N |
| | | | | 17 | 110494125 | *Gat2 l* | N | N | N |
| | | | | 23 | 110503031 | *Gat2 m* | N | N | N |
| | | | | 12 | 110537741 | *Gat2 n* | Brain | N | N |
| | | | | 12 | 110537636 | *GaT2 o* | Y | Y | N |
| | | | | 17 | 110494093 | *Gat2 p* | Y | Y | Y |
| | | | | 17 | 110494094 | *Gat2 q* | Y | Y | Y |
| | | | | 17 | 110494095 | *Gat2 r* | Y | Y | Y |
| $ATB^{0,+}$ | slc6a14 | 1 | C | 7 | 110528352 | *$ATB^{0,+}$* | Y | Y | N |
| $B^0AT2$ | slc6a15 | 4 | N | 1 | 110524453 | *$B^0AT2$ a* | Y | **N** | Y |
| | | | | 2 | 110499724 | *$B^0AT2$ b* | Y | Y | Y |
| | | | | 12 | 118937783 | *$B^0AT2$ c* | Y | Y | **N** |
| | | | | 16 | 110491573 | *$B^0AT2$ d* | Y | Y | Y |
| $B^0AT3$ | slc6a17 | 3 | N | 9 | 110531270 | *$B^0AT3$ a* | Y | N | Y |
| | | | | 16 | 110491175 | *$B^0AT3$ b* | Y | N | Y |
| | | | | 17 | 110494349 | *$B^0AT3$ c* | Y | N | N |
| XT2 | slc6a18 | 2 | N | 32 | 110489359 | *XT2 a* | Y | N | N |
| | | | | 32 | 110489437 | *XT2 b* | Y | N | N |
| $B^0AT1$ | slc6a19 | 6 | N | 2 | 110514444 | *$B^0AT1$ a* | Y | N | N |
| | | | | 3 | 110520772 | *$B^0AT1$ c* | Y | N | N |
| | | | | 2 | 110516599 | *$B^0AT1$ b* | Y | N | N |
| | | | | 3 | 110519055 | *$B^0AT1$ d* | Y | N | N |
| | | | | 18 | 110496758 | *$B^0AT1$ e* | Y | N | **N** |
| | | | | 32 | 110489357 | *$B^0AT1$ f* | Y | **N** | Y |
| SIT1 | slc6a20 | 2 | N | 2 | 110539196 | *SIT1 a* | Y | Y | N |
| | | | | 3 | 110511872 | *SIT1 b* | Y | Y | N |
| Cat1 | slc7a1 | 3 | C | 10* | 110533230* | *Cat1 a** | Y | Y | Y |
| | | | | 10 | 110533232 | *Cat1 b* | Y | Y | Y |
| | | | | 27 | 110507241 | *Cat1 c* | **N** | **N** | **N** |
| | | | | 14 | 110489197 | *Cat2 a* | Y | N | N |

| | | | | | | | | | |
|---|---|---|---|---|---|---|---|---|---|
| Cat2 | slc7a2 | 4 | C | 31 | 110504480 | *Cat2 b* | Y | N | N |
| | | | | 31 | 110504711 | *Cat2 c* | Y | Y | N |
| | | | | Y | 110510086 | *Cat2 d* | **N** | **N** | **N** |
| Cat3 | slc7a3 | 4 | C | 10 | 110534152 | *Cat3 a* | Y | Y | Y |
| | | | | 14 | 110488786 | *Cat3 b* | Y | Y | Y |
| | | | | 31 | 110504989 | *Cat3 c* | **N** | **N** | **N** |
| | | | | Y | 110510151 | *Cat3 d* | Y | Y | Y |
| Cat4 | slc7a4 | 2 | C | 6 | 110525572 | *Cat4 a* | Y | N | N |
| | | | | 11 | 110535941 | *Cat4 b* | Y | **N** | **N** |
| LAT1 | slc7a5 | 5 | N | 1 | 110511826 | *LAT1 a* | Y | Y | Y |
| | | | | 2 | 110493900 | *LAT1 b* | Y | Y | Y |
| | | | | 6 | 110514126 | *LAT1 c* | **N** | **N** | **N** |
| | | | | 9 | 110532220 | *LAT1 d* | Y | Y | Y |
| | | | | 16 | 110491824 | *LAT1 e* | Y | Y | Y |
| $y^+$LAT2 | slc7a6 | 5 | C | 6 | 110526237 | *$y^+$ lat2 a* | **N** | **N** | **N** |
| | | | | 8 | 110530061 | *$y^+$ lat2 b* | Y | Y | Y |
| | | | | 32 | 110487934 | *$y^+$ lat2 c* | Y | Y | Y |
| | | | | 18 | 110496550 | *$y^+$ lat2 d* | N | N | N |
| | | | | 26 | 110506393 | *$y^+$ lat2 e* | **N** | **N** | **N** |
| $y^+$LAT1 | slc7a7 | 2 | C | 9 | 110531642 | *$y^+$ lat1 a* | Y | N | N |
| | | | | 21 | 110500506 | *$y^+$ la12 b* | Y | **N** | **N** |
| LAT2 | slc7a8 | 3 | N | 8 | 110530618 | *LAT2 a* | Y | Y | Y |
| | | | | 9 | 110531623 | *LAT2 b* | Y | Y | Y |
| | | | | 21 | 110500480 | *LAT2 c* | Y | Y | Y |
| $B^{0,+}$AT | slc7a9 | 3 | C | 19 | 110497769 | *$B^{0,+}$ AT a* | N | N | N |
| | | | | 25 | 110505695 | *$B^{0,+}$ AT b* | **N** | **N** | **N** |
| | | | | 26 | 110510393 | *$B^{0,+}$ AT c* | Y | Y | **N** |
| Asc-1 | slc7a10 | 4 | N | 1 | 110524148 | *Asc-1 a* | Y | N | N |
| | | | | 2 | 110499202 | *Asc-1 b* | Y | **N** | **N** |
| | | | | 6 | 110514610 | *Asc-1 c* | Y | Y | Y |
| | | | | 26 | 110516700 | *Asc-1 d* | Y | Y | Y |

| | | | | | | | | | |
|---|---|---|---|---|---|---|---|---|---|
| xCt | slc7a11 | 4 | A | 15 | 110490496 | *xCt a* | Y | Y | N |
| | | | | 21 | 110499969 | *xCt b* | Y | Y | **N** |
| | | | | 14 | 110487732 | *xCt c* | Y | Y | Y |
| | | | | 31 | 110505014 | *xCt d* | Y | Y | Y |
| Asc-2 | slc7a12 | 0 | N | Not Found | | | | | |
| AGT1 | slc7a13 | 0 | N | Not Found | | | | | |
| slc7a14 | slc7a14 | 3 | C | 18 | 110527386 | *slc7a14 a* | Y | **N** | **N** |
| | | | | 7 | 110527387 | *slc7a14 b* | N | N | N |
| | | | | 15 | 110490917 | *slc7a14 c* | Y | Y | **N** |
| MCT7 | slc16a6 | 5 | N | 13 | 110485325 | *MCT7 a* | Y | Y | Y |
| | | | | 13 | 110485326 | *MCT7 b* | Y | Y | Y |
| | | | | 12 | 118937837 | *MCT7 c* | Y | Y | Y |
| | | | | 20 | 110499208 | *MCT7 d* | Y | N | N |
| | | | | 23 | 110502925 | *MCT7 e* | N | N | N |
| TAT1 | slc16a10 | 3 | N | 4 | 110522149 | *TAT1 a* | Y | Y | N |
| | | | | 8 | 110529198 | *TAT1 b* | Y | Y | N |
| | | | | 25 | 110504195 | *TAT1 c* | N | N | N |
| VGLUT2 | slc17a6 | 3 | A | 6 | 110526515 | *VGLUT2 a* | Y | N | N |
| | | | | 26 | 110506645 | *VGLUT2 b* | Y | N | N |
| | | | | 30 | 110521189 | *VGLUT2 c* | Y | N | N |
| VGLUT1 | slc17a7 | 4 | A | 12 | 118937784 | *VGLUT1 a* | Y | N | N |
| | | | | 13 | 110487029 | *VGLUT1 b* | Y | N | N |
| | | | | 16 | 110491576 | *VGLUT1 c* | Y | N | N |
| | | | | 20 | 110491576 | *VGLUT1 d* | Y | N | N |
| VGLUT3 | slc17a8 | 2 | A | 1 | 110522804 | *VGLUT3 a* | Y | Y | N |
| | | | | 2 | 110497771 | *VGLUT3 b* | Y | Y | N |
| Ornt2 | slc25a2 | 0 | C | Not Found | | | | | |
| AGC1 | slc25a12 | 4 | A | 3 | 110520101 | *AGC1 a* | Y | Y | Y |
| | | | | 22 | 110501456 | *AGC1 b* | Y | Y | Y |
| | | | | 8 | 110530664 | *AGC1 c* | Y | Y | Y |
| | | | | 28 | 110509207 | *AGC1 d* | Y | Y | Y |
| AGC2 | slc25a13 | 2 | A | 18 | 110509207 | *AGC2 a* | Y | N | Y |
| | | | | 32 | 110488013 | *AGC2 b* | Y | N | Y |

| | | | | | | | | | |
|---|---|---|---|---|---|---|---|---|---|
| Ornt1 | slc25a15 | 3 | C | 10 | 110534225 | *Ornt1 a* | N | N | N |
| | | | | 27* | 110507247* | *Ornt1 b** | N | N | N |
| | | | | 27 | 110508105 | *Ornt1 c* | Y | Y | Y |
| GC2 | slc25a18 | 0 | A | Not Found | | | | | |
| GC1 | slc25a22 | 7 | A | 1 | 110523386 | *GC1 a* | Y | Y | Y |
| | | | | 2 | 110516669 | *GC1 b* | Y | Y | N |
| | | | | 6 | 110526530 | *GC1 c* | Y | Y | N |
| | | | | 6 | 110518525 | *GC1 d* | Y | Y | N |
| | | | | 7 | 110527759 | *GC1 e* | Y | Y | Y |
| | | | | 17 | 110494701 | *GC1 f* | Y | Y | Y |
| | | | | 30 | 110521172 | *GC1 g* | Y | Y | Y |
| SAMC | slc25a26 | 1 | N | 17 | 110494503 | *SAMC* | Y | Y | Y |
| Ornt3 | slc25a29 | 2 | C | 4 | 110522494 | *Ornt3 a* | N | N | N |
| | | | | 19 | 110498176 | *Ornt3 b* | Y | Y | Y |
| slc25a38 | slc25a38 | 4 | N | 12 | 110538473 | *slc25a38 a* | Y | Y | Y |
| | | | | 13 | 110486514 | *slc25a38 b* | Y | Y | Y |
| | | | | 13 | 110485796 | *slc25a38 c* | Y | Y | Y |
| | | | | 17 | 110493547 | *slc25a38 d* | Y | Y | Y |
| slc25a44 | slc25a44 | 2 | N | 6 | 110526738 | *slc25a44 a* | Y | Y | Y |
| | | | | 26 | 110517597 | *slc25a44 b* | Y | Y | Y |
| VIAAT | slc32a1 | 5 | N | 4 | 110493547 | *VIAAT a* | N | N | N |
| | | | | 7 | 110527704 | *VIAAT b* | Y | N | N |
| | | | | 17 | 110494768 | *VIAAT c* | Y | N | N |
| | | | | 9 | 110532052 | *VIAAT d* | Y | N | N |
| | | | | 16 | 110492348 | *VIAAT e* | Y | N | N |
| PAT1 | slc36a1 | 2 | N | 14 | 110489216 | *PAT1 a* | Y | Y | Y |
| | | | | 31 | 110504544 | *PAT1 b* | Y | Y | Y |
| PAT2 | slc36a2 | 0 | N | Not Found | | | | | |
| PAT3 | slc36a3 | 0 | N | Not Found | | | | | |
| PAT4 | slc36a4 | 1 | N | 27 | 110507984 | *PAT4* | Y | Y | Y |
| SNAT1 | slc38a1 | 0 | N | Not Found | | | | | |
| SNAT2 | slc38a2 | 2 | N | 21 | 110499958 | *SNAT2 a* | Y | Y | Y |
| | | | | 15 | 110490511 | *SNAT2 b* | Y | Y | Y |

| | | | | | | | | | |
|---|---|---|---|---|---|---|---|---|---|
| SNAT3 | slc38a3 | 8 | N | 7 | 110528155 | *SNAT3 a* | Y | Y | Y |
| | | | | 17 | 110494279 | *SNAT3 b* | Y | Y | Y |
| | | | | 7 | 110527782 | *SNAT3 c* | Y | Y | N |
| | | | | 17 | 110494674 | *SNAT3 d* | Y | Y | N |
| | | | | 9 | 110532146 | *SNAT3 e* | N | N | Y |
| | | | | 16 | 110492273 | *SNAT3 f* | N | N | Y |
| | | | | 9 | 110532471 | *SNAT3 g* | Y | N | Y |
| | | | | 16 | 110492139 | *SNAT3 h* | Y | N | Y |
| SNAT4 | slc38a4 | 2 | N | 15 | 110490512 | *SNAT4 a* | Y | Y | Y |
| | | | | 21 | 110499957 | *SNAT4 b* | Y | Y | Y |
| SNAT5 | slc38a5 | 0 | N | Not Found | | | | | |
| SNAT6 | slc38a6 | 2 | N | 19 | 110497839 | *SNAT6 a* | N | N | N |
| | | | | 25 | 110505626 | *SNAT6 b* | Y | Y | Y |
| SNAT7 | slc38a7 | 2 | N | 6 | 110526454 | *SNAT7 a* | Y | Y | Y |
| | | | | 26 | 110506580 | *SNAT7 b* | Y | Y | Y |
| SNAT8 | slc38a8 | 2 | N | 2 | 110495203 | *SNAT8 a* | Y | N | N |
| | | | | 26 | 110506232 | *SNAT8 b* | N | N | Y |
| slc38a9 | slc38a9 | 1 | C | 5 | 110523173 | slc38a9 | Y | Y | Y |
| SNAT10 | slc38a10 | 2 | N | 20 | 110499205 | *SNAT10 a* | Y | N | Y |
| | | | | 23 | 110503021 | *SNAT10 b* | Y | Y | Y |
| slc38a11 | slc38a11 | 1 | N | 3 | 110505257 | *slc38a11* | N | N | Y |
| LAT3 | slc43a1 | 4 | N | 10 | 110534431 | *LAT3 a* | Y | Y | Y |
| | | | | 19 | 110498668 | *LAT3 b* | Y | Y | Y |
| | | | | 14 | 110488829 | *LAT3 c* | Y | Y | Y |
| | | | | 31 | 110505037 | *LAT3 d* | Y | Y | Y |
| LAT4 | slc43a2 | 4 | N | Y | 110517633 | *LAT4 a* | Y | Y | N |
| | | | | 10 | 110534814 | *LAT4 b* | Y | Y | N |
| | | | | 24 | 110503649 | *LAT4 c* | Y | Y | Y |
| | | | | 27 | 110507524 | *LAT4 d* | Y | Y | Y |
| SFXN1 | SLC56A1 | 4 | N | 1 | 110530947 | *SFXN1 a* | Y | Y | Y |
| | | | | 14 | 110487787 | *SFXN1 b* | Y | Y | Y |
| | | | | 23 | 110502652 | *SFXN1 c* | Y | Y | N |
| | | | | 31 | 110504533 | *SFXN1 d* | Y | Y | Y |

| | | | | | | | | | |
|---|---|---|---|---|---|---|---|---|---|
| SFXN2 | slc56a2 | 1 | N | 23 | 110502746 | *SFXN2* | Y | Y | Y |
| SFXN3 | slc56a3 | 0 | N | Not Found | | | | | |
| Cystinosin | CTNS | 2 | N | 1 | 110503198 | *CTNS a* | Y | Y | Y |
| | | | | 5 | 110524220 | *CTNS b* | Y | Y | Y |
| pqlc2 | slc66a1 | 2 | N | 9 | 110532347 | *pqlc2 a* | Y | Y | Y |
| | | | | 16 | 110492259 | *pqlc2 b* | Y | Y | Y |

* : for *Cat1a* and *Ornt1b* , geneID and chromosom position correspond to Omyk_1.0 genome

All CAAT refered in this table were previously identified by Morin and coworkers (https://doi.org/10.1016/j.isci.2024.108894)
