## Supplementary Table 2 for "Dissecting the nutritional regulations of a whole amino acid transporter family from a complex genome species: A holistic approach turning weaknesses into strengths"

**Table of newly designed and validated primers**

| Genes | Forward Primer | Reverse Primer | Efficiency |
|---|---|---|---|
| EAAT3 a & EAAT3 b | 5'-GATCTCTTCCAGCTCTGCCA-3' | 5'-GAGCTGGGCAATGAAGATGG-3' | 1,97 |
| EAAT2 a | 5'-AGGCCCAGGAACAAGTGTAT-3' | 5'-GAACGCAATCACCATGACGA-3' | 1,87 |
| EAAT2 b | 5'-ATGCTGGTCAGATCGTCACT-3' | 5'-GTATGAGTCTCCCACCACGT-3' | 1,93 |
| EAAT1 a & EAAT1 c | 5'-TCCTGAGACGCAACCTCTTT-3' | 5'-CCATCTTCCCAGACGCTTTG-3' | 1,98 |
| EAAT1 b & EAAT1 d | 5'-ATCGTGGTGCTCATCATCCA-3' | 5'-CTCAGTCAGGTTGGTCAGGT-3' | 2,02 |
| EAAT1 e | 5'-TATGACCAAGAGCAACGGGG-3' | 5'-AGTGCAAATCCCAGCCCTAT-3' | 1,92 |
| ASCT1 a & ASCT1 b | 5'-GGTCATCCTGCCTCTCATCT-3' | 5'-GGATGAAACGACTGATCCGC-3' | 2,00 |
| ASCT2 a & ASCT2 b | 5'-ATGGCGTTCATCATCTCCCC-3' | 5'-GAAAGGCAGCAGACACCAAG-3' | 1,93 |
| EAAT6 a & EAAT6 b | 5'-CCTCCTCCCCTTCTTCTACC-3' | 5'-CCGATCCACATTGCAGTTCT-3' | 2,04 |
| EAAT7 a & EAAT7 b | 5'-AGCCATCTTCATCGCTCAGA-3' | 5'-GGTACGCATTCTGTCAAGCA-3' | 1,80 |
| Gat1 a | 5'-GACAGCACCACTCACCTTTC-3' | 5'-GGAAGGTTAGATGGGCAGGA-3' | 2,08 |
| Gat1 b | 5'-TCCATGCTCCTGATACCTGG-3' | 5'-TCGGAGACAGGCTGAATAGAA-3' | 1,80 |
| Gat1 d | 5'-TGTTCAGTGCAGTCCAAATGG-3' | 5'-GCAATGTTGGTCTCGGCAA-3' | 1,95 |
| Gat1 e | 5'-CTCTGTTGGTTGTTGGGGTC-3' | 5'-GGCACACAGACACATCATGG-3' | 2,04 |
| GlyT2 b | 5'-AGACACATACGCTGCCTCTT-3' | 5'-TAGGAGTCACAAAGGCCCAG-3' | 1,99 |
| TAUT a & TAUT c | 5'-GGGGTCTGGGTCATCTGTTT-3' | 5'-ACGTTGCAGGTCTGGGTAC-3' | 1,98 |
| TAUT b & TAUT d | 5'-TATCACTCCCGTCCTCTGTG-3' | 5'-AACTCGCACCTCCCTTTACT-3' | 1,97 |
| GlyT1 a & GlyT1 b | 5'-ACAGAAATGGAGGAGGAGCC-3' | 5'-CCACACCTTTGAACATGGGG-3' | 2,03 |
| Gat2 a & Gat2 b | 5'-GCCACAGAGCGATGAGAAGA-3' | 5'-AGGGGATGAAGAAAGCACCT-3' | 2,01 |
| Gat2 c | 5'-GCTTTAGTGGCCTTCATCTCC-3' | 5'-ATGTGGCTGGGGTGACATAC-3' | 1,91 |
| Gat2 d & Gat2 e | 5'-TGTCCCATGTTCGAAGGAGT-3' | 5'-GGAGAGGTGTGGGTTCAACA-3' | 1,98 |
| Gat2 i | 5'-GTTCGGAGGTCTGGGCTTT-3' | 5'-ACAGTCAAACAACACACAGGT-3' | 1,90 |
| Gat2 k | 5'-AGGTTTGGATGGATGCAGGA-3' | 5'-ATGTCCACACCCTGTTCTCC-3' | 1,97 |
| Gat2 n | 5'-GTGTGCACAGCTACCTTCATC-3' | 5'-AGGGCCATCTGTTCTGCAT-3' | 1,88 |
| Gat2 o | 5'-TCACCACTACTTCTGCTGCC-3' | 5'-TCCTGAGTACCTGATTCCTTTCC- | 2,01 |

| | | | |
|---|---|---|---|
| Gat2 p & Gat2 q & Gat: | 5'-GTTCACTCTCTCCTCCACCC-3' | 5'-CCGTGTCTCTGTCTAGGAGG-3' | 1,93 |
| Gat2 f | 5'-GAGGGTATCGGCTATGCAGG-3' | 5'-AGAGGAGAAGTCAACACAGTCA-3 | 1,95 |
| $B^0$AT2 a | 5'-ACGAGTGATGATGTCTCCCC-3' | 5'-GAGATCCGGGTAGTGTCTGC-3' | 1,95 |
| $B^0$AT2 c | 5'-ACAACCCCATCCACAACAAC-3' | 5'-TGTCGTTTAACAGGCCCAGA-3' | 2,02 |
| B°AT2 d | 5'-GCGAGGATGGTCAGGAGG-3' | 5'-AGCTCCAGGAAGAACAAGGG-3' | 1,97 |
| $B^0$AT2 b | 5'-TGTCTGCCCCGTGTATGTG-3' | 5'-GTTCTGAGGGCTTAAAGGTGG-3' | 2,02 |
| $B^0$AT3 a | 5'-GCACAGAGGAGAGATGGGTG-3' | 5'-CGGGTCAGAGGTCAGAGATG-3' | 2,09 |
| $B^0$AT3 c | 5'-GAAGATTGTTCTCGAGCCCC-3' | 5'-CGCTAGCTCCAGGAAGAAGA-3' | 1,89 |
| $B^0$AT3 b | 5'-CTCTCCCATGCCTTCTCACC-3' | 5'-GGGTTGTGTTGTGTTGGGTT-3' | 1,95 |
| XT2 a & XT2 b | 5'-ACGGAATAGGACGGTTCAGT-3' | 5'-CTCTGTGTCGGGGAAGTTCA-3' | 1,98 |
| $B^0$AT1 a & B0AT1 b | 5'-CATCTACACCCGAGGAACCA-3' | 5'-GGAAAGGAATCATGAATGCACCT | 1,98 |
| $B^0$AT1 c & B0AT1 d | 5'-CCCCGCCATCATGTTTGTC-3' | 5'-ACTCATTATCCTTCTTGCAGCAG- | 1,98 |
| $B^0$AT1 e | 5'-CCTACCTCGTATGGGACCAG-3' | 5'-CCTGTTACTTTTAGTCCATCGGT- | 2,02 |
| $B^0$AT1 f | 5'-GGAGGCGATCACTGGTATTG-3' | 5'-CCAGAACAGGTTGGGCTTG-3' | 1,94 |
| SIT1 a & SIT1 b | 5'-TGTCCTGGCGTGTGTATCAT-3' | 5'-CTCTCCATGCTCCAATGCTG-3' | 2,00 |
| LAT1 a & LAT1 b | 5'-TACGCCTTCTCCAACGACAT-3' | 5'-GATAGGCAGCAGGATATTCACC-3 | 1,93 |
| LAT1 d & LAT1 e | 5'-GGAGGCCCATCAAGATTTGG-3' | 5'-GTACCCCAGAAGATAGGCGG-3' | 1,88 |
| LAT2 a | 5'-GCCACAATGCTTCAACACCT-3' | 5'-TGGGTTGAACACATCTCCTAGA-3 | 1,90 |
| LAT2 b & LAT2 c | 5'-GGCCTTCCTGCTCATCTTCT-3' | 5'-CCCTTCGTTTCCCCTGGATA-3' | 2,05 |
| Asc-1 a | 5'-TGGAAGAAGCCCAAACTTTACA-3' | 5'-AGGTTGCTGACTCGATGAAGT-3' | 2,00 |
| Asc-1 b | 5'-TGCAGACATGGATGGAGGAA-3' | 5'-ACTGAACCCGAGTGCTCTAG-3' | 1,93 |
| Asc-1 c & Asc-1 d | 5'-TCGCCGTCAGATAGAGTCAC-3' | 5'-CAGCTCAGCGTAACACATGG-3' | 1,96 |
| xCt a & xCt b | 5'-GCTGATCAACTTTGCCTCCT-3' | 5'-GGTGCACGGTCAGGTAGTA-3' | 2,01 |
| xCt c & xCt d | 5'-TCTGGGAGTGTTGGAATGTCT-3' | 5'-CATCTGTGGTCCAAAGGCCT-3' | 1,93 |
| MCT7 a & MCT7 b & M | 5'-TGTCCCAGTACTTCAGCAGG-3' | 5'-CTGGTCAGGTCGGATGATGA-3' | 1,85 |
| MCT7 d | 5'-TCCATCTGCGTCTTCACTATGA-3' | 5'-GGCAGGAAGGTCAGACAGTA-3' | 1,91 |
| TAT1 a & TAT1 b | 5'-GAGCAAGGTGTTCAACGTCA-3' | 5'-CACATGAGCAGAACCTCCTTG-3' | 1,89 |
| VGLUT2 a & VGLUT2 | 5'-ATGGAGTCCGTCAAGATGAGA-3' | 5'-ACAGAGGTGCTTTCTTCTCCT-3' | 1,92 |

| | | | |
|---|---|---|---|
| VGLUT2 c | 5'-ATCCCGAACCCGTTAAGCAT-3' | 5'-GCAGTCACATAAAGGCGGTT-3' | 2,13 |
| VGLUT1 a & VGLUT1 | 5'-TGCTCGTACACACTACAGCT-3' | 5'-CCCAAAACTGCCATAGACGT-3' | 1,91 |
| VGLUT1 c & VGLUT1 | 5'-GGCAGCACGAACTCACATT-3' | 5'-GCATAGGACCCACAAAAGGC-3' | 2,01 |
| VGLUT3 a & VGLUT3 | 5'-CGCAGTCGTAAAATCATGTCAAC- | 5'-GAACCCTGAGATGGCAAAGC-3' | 1,82 |
| AGC1 a & AGC1 b | 5'-CTCCACCAGCTCCTTCGTAG-3' | 5'-CCTCTGTGGTGAATTTGTCTCTT- | 1,89 |
| AGC1 c & AGC1 d | 5'-CAGGGTGATGGTTCCAGGT-3' | 5'-TTCTCCTACGAACGAGCCTG-3' | 1,96 |
| AGC2 a & AGC2 b | 5'-CTGGGAGCAGTAGAGTCCTG-3' | 5'-TCCTGTCAATGTCCACCAGA-3' | 1,95 |
| GC1 a | 5'-ATCGGAGGAGGAGGAAGGAA-3' | 5'-GTCAAACAGTCCAGCATCCC-3' | 1,97 |
| GC1 b | 5'-ACATATCAACTCTCTGACGACCA- | 5'-CCCTCTGTACATCCCGAAGT-3' | 1,82 |
| GC1 c & GC1 d | 5'-GGTCAGAAGCTTAGCCTGC-3' | 5'-TGCTGTATGGGACCTCAACT-3' | 2,04 |
| GC1 e | 5'-TTTCCATTGTCTCGCGTTCG-3' | 5'-TCTGCTGCTTGGTCATGGTA-3' | 1,93 |
| GC1 f | 5'-AACGAGGAGAGCTACAACGG-3' | 5'-GCTGGTCTTGTCTCATGCAG-3' | 1,97 |
| GC1 g | 5'-AGGATCCAGTCTATGAACAGGG-3 | 5'-TGTTCTTGTCACTTAGTTGCCA-3' | 2,01 |
| SAMC | 5'-GAATAGCAGCGTTCGTCACC-3' | 5'-GTGCCAGAAAACAGACCCG-3' | 2,05 |
| slc25a38 a & slc25a38 | 5'-CCCACATCTGTACAGGAGGA-3' | 5'-TCAGGACTTGAGCCCCATG-3' | 1,85 |
| slc25a38 c & slc25a38 | 5'-GGTCAAGACTCACATCCAGGT-3' | 5'-ATCAGTTCCTCCCTGTCTCC-3' | 1,98 |
| slc25a44 a & slc25a44 | 5'-GGATATGTGGCCTCCCTTCT-3' | 5'-TGCCTTCAACCTGCACTCTA-3' | 1,94 |
| VIAAT b & VIAAT e | 5'-TCTAAAGCCTCAGGTCTGGG-3' | 5'-GTCTTCGTCCTCCTCGTACA-3' | 1,82 |
| VIAAT c & VIAAT d | 5'-GTGTAACGAGTTGGCATCAGA-3' | 5'-GCGCACCAGAATTCCATCC-3' | 1,96 |
| PAT1 a & PAT1 b | 5'-GGCATCGTCACCTTCCTCTA-3' | 5'-TGAAGGGCAAAGGTGATGTAGA- | 2,09 |
| PAT4 | 5'-CACCAGGGCATCACGTTTAC-3' | 5'-GTGAACACAGACCACACCC-3' | 2,00 |
| SNAT2 a & SNAT2 b | 5'-GATTATCATCCAGGCACCGC-3' | 5'-AGGGAGAAGATGGACACAGC-3' | 1,94 |
| SANAT3 a & SNAT3 b | 5'-TCAGTGTCATCCTGCCTCTG-3' | 5'-TAGGCCGTCTGTGAGTTGAG-3' | 1,94 |
| SNAT3 c & SNAT3 d | 5'-ATCTACACTGAGCTCCGCAA-3' | 5'-GGGAAGAGCACAATGGGGA-3' | 1,83 |
| SNAT3 e & SNAT3 f | 5'-CCTACAGCAAAGTGGACCCT-3' | 5'-CACGAAGAGGAGACACAGGG-3' | 1,96 |
| SNAT3 g & SNAT3 h | 5'-CTGGCTCCATCATCACCATG-3' | 5'-GTGGAAAGATGATGATGACGCT- | 1,91 |
| SNAT6 b | 5'-CCTGGCTTTCTCCTTCCTCT-3' | 5'-CTCTGAGTCTACGTGGCTGT-3' | 1,92 |
| SNAT7 a & SNAT7 b | 5'-TGTTGGCACCTGGTATGTCA-3' | 5'-CACACTGCTCACATGACACT-3' | 1,90 |

| | | | |
|---|---|---|---|
| SNAT8 a | 5'-TCATCTTCCCAGGGCTATGTC-3' | 5'-ACTCCCCTATTCCTACCCAGT-3' | 2,07 |
| SNAT8 b | 5'-CAAAGTCAGGGTGGTGTGC-3' | 5'-AGTGCTGGTGATTTTGGTTGT-3' | 2,00 |
| SNAT10 a | 5'-CCCCAGATCAAAGGACCTGA-3' | 5'-CTCAGACTGCCCCTCTTCTC-3' | 1,97 |
| SNAT10 b | 5'-CCCCAGATCAAAGGACCTGA-3' | 5'-CTCAGACTCTTCTCCTCCCG -3' | 2,09 |
| slc38a11 | 5'-GGAGAATGGCGAAGGGAAAC-3' | 5'-TGCTACCAAAACCAGGAGAAAA-3 | 1,91 |
| LAT3 a & LAT3 b | 5'-ATGAAGGAGTGCGACGTAGA-3' | 5'-GGTGTGGACGATGAAGGAAATT-: | 1,92 |
| LAT3 c | 5'-ACATTCTGTCAACGTTGTGTCA-3' | 5'-ATCGTGTCGTTCTCTGAGCA-3' | 1,82 |
| LAT3 d | 5'-TGTGGCAGAGACAGAGAAGAG-3' | 5'-GTGGAGGATGAAGGTCAGGA-3' | 1,97 |
| LAT4 b | 5'-CAACATCCAACTGAACTCCTGT-3' | 5'-TTCAGAGAACGGGGAGAGC-3' | 1,85 |
| LAT4 a | 5'-CTCTGAACTTCTGTCTCGGC-3' | 5'-GTCAACGCCATCCACCATC-3' | 1,94 |
| LAT4 c & LAT3 d | 5'-GTACCTCCAGATGAACGGCT-3' | 5'-TGGACAGATTCTTGGGGTCG-3' | 1,95 |
| SFXN1 a | 5'-AGACTGGCACCGTGTACATC-3' | 5'-CGGCTCCTTGATGTTAATGTCC-3 | 1,96 |
| SFXN1 c | 5'-AAGCGGTTCCCAGTCCTAAA-3' | 5'-GGGATTATGCATGGTGGGTG-3' | 2,00 |
| SFXN1 b & SFXN1 d | 5'-TGCTCCTCTCTCAGTCAATCA-3' | 5'-AATGCCGTGTTTCAGTTCCC-3' | 1,78 |
| SFXN2 | 5'-TCACGACTGGATGAGGCTAA-3' | 5'-ACCCACTGCCAGAACACTAC-3' | 1,96 |
| CTNS a | 5'-GGCAGCTTCAGTCTCATTCAG-3' | 5'-TAGGAAGTCATTTGGCCCCA-3' | 1,98 |
| CTNS b | 5'-GGAGGCAGCTTCAGTCTCAT-3' | 5'-AATCGTTTGGACTCACTGGC-3' | 2,07 |
| ATF4 a | 5'-GCTGTCCTTCCCTTGGCTAT-3' | 5'-GGGCTGCAGGAACCAATAAG-3' | 1,93 |
| ATF4 b | 5'-AATTCTCCTTCTATCCCCTCCTC-3 | 5'-GGTCAAACTCACTCAGGTCG-3' | 1,91 |
| ATF4 c & ATF4 d | 5'-TTGTTTAGTTTGTCTCCTTCCCC-3 | 5'-CTTCTCAGCCATCCAGTCCA-3' | 1,80 |

**Table of primers previously published**

| Genes | Forward Primer | Reverse Primer | Refs |
|---|---|---|---|
| 4f2hc b | 5’-GGAGAGTGGTCGGAACAAGT-3’ | 5’-TCTCCTGGTCCAACTCGTTC-3’ | |
| 4f2hc d | 5’-CACCTCGCTGCAAAGACATT-3’ | 5’-TTGCCAACGTCAGAGGAGAT-3’ | |
| Cat1 a & Cat1 b | 5’-GCTGCACTGTCAACATCACT-3’ | 5’-TGGTCTTGCTCTCTGGTTGT-3’ | |
| Cat3 a | 5’-GCGAACGTGAAGTCAACTGT-3’ | 5’-CAACAAGATCCGCCCAAAGG-3’ | |
| Cat3 b | 5’-CGGACTGTAAAGTTCAATCGGA-3’ | 5’-GGAGCAGCATCTTACCGAAG-3’ | |

| | | | |
|---|---|---|---|
| Cat3 d | 5'-ACACCAACACCTCCAGCATA-3' | 5'-CCCGAAGTAGGCGAAGAAAC-3' | https://doi.org/10.1016/j.isci.2024.108894 |
| y+LAT2 b | 5'-CCATCTCCATGCCCATTGTG-3' | 5'-AGGCGTTGAGTCCTCCATAG-3' | |
| y+LAT2 c | 5'-GTATCGGGGTGGCTCTTTCT-3' | 5'-TCCAGGTCTAGGTCAGGTGA-3' | |
| Ornt1 c | 5'-ACGCTGTCCTTTTCATGAGC-3' | 5'-GGACCACACCGAACTCTTCT-3' | |
| Ornt3 b | 5'-ACATTGGACTTTGCAGCAGG-3' | 5'-GGCGTTGATGAAGGTCAGAC-3' | |
| SNAT4 a & SNAT4 b | 5'-TCCTTTGGTATGTCGGTCTTCA-3' | 5'-GCAGTCATTAGCAGCAAGTGA-3' | |
| slc38a9 | 5'-GTATTGACATAGTGCCTGCGG-3' | 5'-GAGTCGGCTGTAGTAGTGCA-3' | |
| Pqlc2 a | 5'-AGCAGTGTTTTGAGAGCGTG-3' | 5'-TCCCAAACCCATACCGATCC-3' | |
| Pqlc2 b | 5'-TTTGGTTTCTGCTGCTGTGG-3' | 5'-GCAGAGAAGCCCAGGACATA-3' | |
| $B^{0,+}$AT c | 5'-GCCTACGATGGGTGGAATCA-3' | 5'-CAAAAGTCACAGCAACAGCG-3' | |
| rBAT | 5'-GAGGAGAGGCAGCAGTGTTA-3' | 5'-AGGATCTTGTTGTGGGTCCA-3' | |
| chop | 5'-CGACAATGTCCAACAACCTG-3' | 5'-ACGAGGAGAACGAGGTGCTA-3' | https://doi.org/10.1080/15548627.2015.1117732 |
| asns | 5'-CTGCACACGGTCTGGAGCTG-3' | 5'-GGATCTCGTCTGGGATCAGGTT-3' | https://doi.org/10.1007/s00726-010-0533-3 |
| EF1α | 5'-TCCTCTTGGTCGTTTCGCTG-3' | 5'-ACCCGAGGGACATCCTGTG-3' | https://doi.org/10.3390/cells9081754 |
